# A CpG island-encoded mechanism protects genes from premature transcription termination

**DOI:** 10.1101/2022.03.24.485638

**Authors:** Amy L. Hughes, Aleksander T. Szczurek, Jessica R. Kelley, Anna Lastuvkova, Anne H. Turberfield, Emilia Dimitrova, Neil P. Blackledge, Robert J. Klose

## Abstract

Transcription must be highly controlled to regulate gene expression and development. However, our understanding of the molecular mechanisms that influence transcription and how these are coordinated in cells to ensure normal gene expression remains rudimentary. Here, we reveal that actively transcribed CpG island-associated gene promoters recruit SET1 chromatin modifying complexes to enable gene expression. Counterintuitively, this effect is independent of SET1 complex histone modifying activity, and instead relies on the capacity of these complexes to interact with the RNA Polymerase II-binding protein, WDR82. Unexpectedly, we discover that SET1 complexes sustain gene transcription by counteracting the activity of the ZC3H4/WDR82 protein complex, which we show can pervasively terminate both genic and extragenic transcription. Therefore, we discover a new gene regulatory mechanism whereby CpG island elements nucleate a protein complex that protects genic transcription from premature termination, effectively distinguishing genic from non-genic transcription to enable gene expression.

## Introduction

Precise control of gene expression is essential for cell viability and normal development. At the most basic level, gene expression is controlled by transcription factors that recognise specific DNA sequences in gene regulatory elements and shape how RNA Polymerase II (RNA Pol II) initiates transcription from the core gene promoter^1^. Beyond these DNA sequence-encoded mechanisms, transcription is also influenced by chromatin or epigenetic states at gene promoters and by mechanisms that regulate elongation of RNA Pol II (reviewed in^1-6^). However, we understand far less about how these additional influences on transcription are coordinated with initiation to control gene expression.

CpG islands (CGIs) are associated with most vertebrate gene promoters and are epigenetically distinct from the rest of the genome in that they evade DNA methylation^7, 8^. Non-methylated CpG dinucleotides in CGIs act as binding sites for CGI-binding proteins, many of which are able to post-translationally modify histones and create distinct chromatin states at gene promoters to regulate gene expression^9^. For example, transcribed CGI-associated gene promoters are enriched for histone H3 lysine 4 trimethylation (H3K4me3) which has been proposed to support gene expression. In this context, H3K4me3 is deposited primarily by SET1 complexes, which contain the SET1A or SET1B histone methyltransferases and a series of shared interaction partners that regulate their chromatin binding and enzymatic activity^10-19^. Importantly, SET1 complexes can recognise non-methylated DNA in CGIs via their CFP1 component, but rely on additional multivalent interactions with CGI chromatin and the transcriptional machinery to localise specifically to actively transcribed genes^19-24^. Once deposited, H3K4me3 is bound by additional reader proteins that have been proposed to further modify histones, remodel nucleosomes, and also directly influence RNA Pol II activity to support gene expression^17, 25-32^.

Based on their capacity to deposit H3K4me3 at active genes, SET1 complexes are generally considered transcriptional activators. However, counterintuitively, when SET1 complexes are disrupted, this often causes both increases and decreases in gene expression, which do not necessarily correlate with alterations in H3K4me3 at affected genes^12, 17, 19, 20, 33-42^. Furthermore, cells lacking the methyltransferase domain of SET1A are viable, yet complete removal of SET1A causes cell and early embryonic lethality, suggesting that SET1 complexes may also regulate gene expression independently of histone methylation^11, 39, 43^. In line with this possibility, SET1 complexes contain a protein called WDR82 that directly interacts with RNA Pol II and this could provide an alternative mechanism to influence RNA Pol II activity and transcription^18, 44, 45^. Nevertheless, despite decades of intense study on SET1 complexes and emerging evidence implicating them in human disease^46-49^, how they actually regulate gene expression remains unclear.

Additional systems that function independently of chromatin to regulate RNA Pol II elongation also have central roles in controlling transcription and gene expression^2, 3^. For example, at most genes there is a checkpoint ∼30-50 base pairs downstream of the transcription start site where initiated RNA Pol II pauses. Pausing is overcome by mechanisms that promote the release of RNA Pol II into productive elongation^3^. However, a large fraction of paused RNA Pol II does not continue into productive elongation and instead undergoes premature transcription termination (PTT)^5, 50-53^. PTT can also occur further into transcribed genes where it is often associated with stable nucleosomes at the boundary of promoter-associated CGIs, or even further into the gene at cryptic polyadenylation sites^5, 54-57^. Transcripts arising from PTT events are then usually subject to rapid turnover by the nuclear exosome^58^. Numerous factors can contribute to PTT, including the Integrator complex which binds to RNA Pol II and instigates PTT by cleaving nascent RNA as it exits the RNA Pol II active site, and the cleavage and polyadenylation (CPA) complex that recognises cryptic polyadenylation signals to promote PTT further into the gene^54, 55, 57, 59-67^. Recently an additional factor, ZC3H4, was shown to attenuate extragenic and long non-coding RNA transcription, resulting in transcription termination^68, 69^. Importantly, ZC3H4 binds to the RNA Pol II-interacting protein WDR82, and binding of ZC3H4 to WDR82 appears to be important for its effects on transcription^13, 45, 68, 70^.

If uncontrolled, PTT could be highly detrimental to gene expression. Therefore, mechanisms have evolved to oppose and regulate this process^4, 5^. For example, in addition to its role as a component of the spliceosome, the U1 snRNP can independently bind to 5’ splice sites in nascent RNA and inhibit the activity of the CPA machinery at downstream cryptic polyadenylation sites ^54, 55, 71-73^. Furthermore, TFIIS can help to restart backtracked RNA Pol II to limit PTT, while SCAF4/8 interact with elongating RNA Pol II and suppress gene-intrinsic polyadenylation site usage ^74, 75^. These examples demonstrate that control of PTT can provide an additional mechanism to regulate gene expression. However, our current understanding of the factors that regulate PTT and the molecular logic underpinning how these systems control gene expression remains rudimentary and is a major conceptual gap in our understanding of gene regulation.

Here we set out to understand how the CGI-binding and chromatin-modifying SET1 complexes regulate gene expression. Using genome engineering, degron approaches, and quantitative genomics we discover that SET1 complexes play a remarkably widespread and highly specific role in enabling normal gene expression. This effect does not rely on their histone methyltransferase activity, nor is it related to effects on H3K4me3, but instead SET1 complexes regulate gene expression by interacting with the RNA Pol II-binding protein WDR82. Unexpectedly, this enables SET1 complexes to support genic transcription from CGI-associated gene promoters by counteracting an opposing ZC3H4/WDR82-dependent PTT activity that we show can target both genic and extragenic transcription. As such, we uncover a new gene regulatory mechanism whereby CGIs nucleate a protein complex which counteracts PTT to distinguish genic transcription from non-genic transcription and enable gene expression.

## Results

### SET1 complexes primarily support gene expression

Despite their intimate association with actively transcribed CGI-associated gene promoters, it remains very poorly understood whether SET1 complexes play a significant role in regulating gene expression^17^. Addressing this important question has been extremely challenging given that SET1 complexes are essential for cell viability and traditional perturbation approaches previously used to study their function are slow. This means existing gene expression analyses will be confounded by pleiotropic secondary effects that inevitably result from deteriorating cell viability. To overcome this limitation we used a rapid degron approach and quantitative time-resolved genomics to examine how SET1 complexes regulate gene expression in mouse embryonic stem cells (ESCs)^76, 77^. We focussed initially on SET1A as it is essential in ESCs and is proposed to contribute centrally to H3K4me3 deposition at actively transcribed genes^11, 12^. We used CRISPR/Cas9-mediated genome engineering to introduce a degradation tag (dTAG) into the SET1A gene^76^. The addition of the dTAG did not affect SET1A protein levels, SET1A complex formation, or its localisation to CGI-associated gene promoters (Fig.1a and Extended Data Fig.1a-e). Importantly, within 2 hours of treatment with dTAG13, there was a near-complete loss of SET1A protein and its occupancy at target genes as assessed by western blot and calibrated chromatin immunoprecipitation coupled to sequencing (cChIP-seq) (Fig.1a and Extended Data Fig.1d-e).

Having shown that we can rapidly deplete SET1A, we then examined gene expression using calibrated total RNA-seq (cRNA-seq) at several time points after SET1A removal (Fig.1b and Extended Data Fig.1g). Remarkably, after only two hours of SET1A depletion we observed profound and widespread effects on gene expression and, in contrast to previous findings^12, 17, 19, 20, 33-39^, we found that the removal of SET1A predominantly resulted in reductions in gene expression (2299 genes) with a much smaller number of genes increasing in expression (414). Importantly, these effects were due to depletion of SET1A, as treating wild type cells with dTAG13 caused no significant alterations in gene expression (Extended Data Fig.1f). Interestingly, changes in gene expression were less pronounced at later time points after SET1A depletion, suggesting that additional mechanisms may compensate for its depletion over time (Extended Data Fig.1g). This highlights the importance of using rapid depletion to capture primary effects on gene expression when studying essential proteins.

Although SET1A is highly expressed and thought to predominate in forming the SET1 complex in ESCs, its paralogue SET1B is also expressed (Fig.1c). Furthermore, whilst the majority of genes were reduced in expression following depletion of SET1A, SET1B expression was increased (Extended Data Fig.1j). This suggested that SET1B might also regulate gene expression and possibly have a compensatory role after SET1A depletion. To examine this possibility and to ensure we had removed all SET1 complex activity, we created cell lines in which we introduced a dTAG into the SET1B gene, and into both SET1A and SET1B (Fig.1c and e). In contrast to SET1A depletion, after 2 hours of SET1B depletion there were only minimal effects on gene expression, which were almost completely absent at later time points (Fig.1d and Extended Data Fig.1h). However, when SET1A and SET1B were removed simultaneously, there was an even more pronounced reduction in gene expression than was observed for depletion of SET1A alone (Fig.1f and Extended Data Fig.1i). Now, 2928 genes were significantly reduced in expression, whereas far fewer genes (745) were increased in expression and these effects were much smaller in magnitude. This suggests that SET1A and SET1B both contribute to gene regulation. This was also evident when we examined the overlap in the genes that rely on SET1 proteins for their expression and it was clear that reductions in expression were largest in magnitude when SET1A and SET1B were simultaneously depleted (Fig. 1g-h). Interestingly, there was no clear enrichment for specific ontology terms amongst genes with reduced expression after SET1A/B depletion, suggesting that there was no defined type of gene that relies on SET1A/B for expression (Extended data Fig.1l). Instead, the only obvious feature shared amongst these genes was that they were more moderately expressed compared to unaffected genes (Fig.1i). Therefore, our rapid depletion approaches reveal that SET1A and SET1B play a prominent and overlapping role in supporting gene expression.

**Fig.1:**
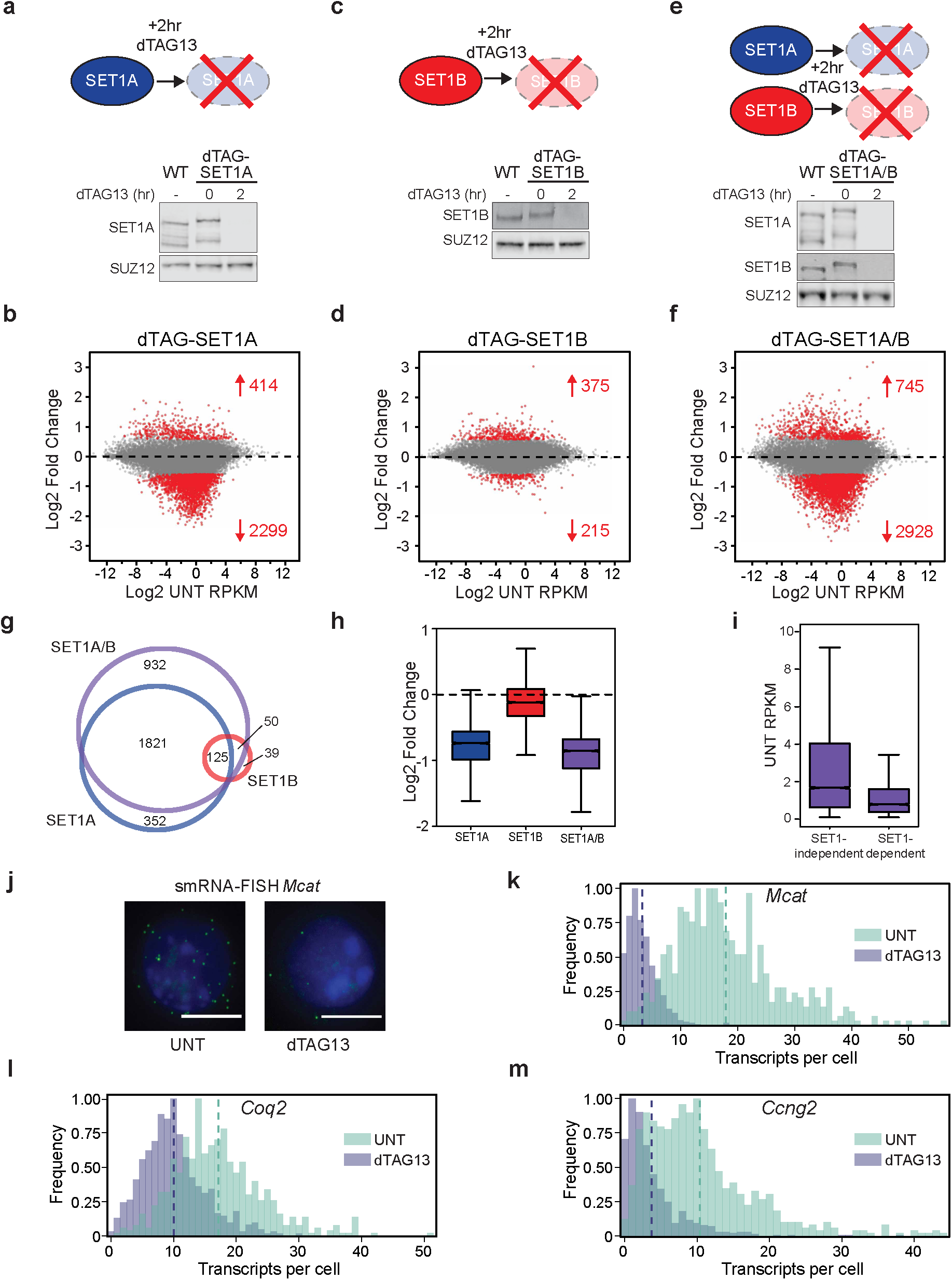
SET1 complexes primarily support gene expression. **(a)** A schematic illustrating the approach used to rapidly deplete SET1A (top panel). A western blot showing comparable SET1A levels in wild type (WT) and dTAG-SET1A lines (bottom panel), and that 2 hours of dTAG13 treatment causes depletion of dTAG-SET1A. SUZ12 functions as a loading control. **(b)** An MA-plot showing log2 fold changes in cRNA-seq signal in the dTAG-SET1A line following 2 hours of dTAG13 treatment (n=20633). Significant gene expression changes (p-adj<0.05 and >1.5-fold) are coloured red and the numbers of significantly changed genes are indicated. **(c-d)** As per a-b but for the dTAG-SET1B line. **(e-f)** As per a-b but for the dTAG-SET1A/B line. **(g)** A Venn diagram showing the overlap between genes with a significant reduction in expression in the dTAG-SET1A, dTAG-SET1B, and dTAG-SET1A/B lines. **(h)** A box plot showing the log2 fold changes in cRNA-seq signal in each of the dTAG cell lines for the complete set of genes that rely on SET1 complexes for expression (n=3320). **(i)** A box plot showing the expression (RPKM) level in untreated cells (UNT) for expressed genes that rely on SET1A/B (SET1-dependent, n=2544) compared to Unchanged genes (SET1-independent, n=9989). **(j)** Example images of smRNA-FISH for the *Mcat* gene in the dTAG-SET1A/B line, showing an untreated (UNT) cell and a cell treated with dTAG13 for 2 hours. Green corresponds to *Mcat* RNAs and blue corresponds to DAPI staining of DNA. The white scale bars correspond to 10 μm. **(k)** A histogram illustrating the number of transcripts per cell as measured by smRNA-FISH before (light green) and after 2 hours of dTAG13 treatment (light purple) for the *Mcat* gene in the dTAG-SET1A/B line. The dashed lines correspond to the mean of the distribution. **(l-m)** As per (k) but for the (l) *Coq2* and (m) *Ccng2* genes.

cRNA-seq analysis measures average changes in gene expression across millions of cells and therefore does not capture how these effects ultimately manifest in individual cells within the population. To address this important question we carried out single molecule RNA fluorescent in situ hybridisation (smRNA-FISH) to enable absolute quantification of gene expression changes in single cells for three SET1-dependent genes (Fig.1j)^78^. Importantly, for each of the genes examined, the reductions in gene expression after SET1A/B depletion were on average uniform across the cell population (Fig.1k-m). Therefore, our genomic and imaging analysis reveal that SET1 complexes have a widespread, overlapping, and uniform role in enabling gene expression.

### SET1 complexes can regulate gene expression independently of H3K4me3 and methyltransferase activity

SET1 complexes are thought to be the predominant H3K4 tri-methyltransferases in animals and the deposition of H3K4me3 has been proposed to influence gene expression^11, 12, 17, 79-81^. As our degron approach allows us to capture the earliest and most primary influences of SET1 protein depletion on gene expression, we set out to examine whether the observed gene expression changes might result from effects on H3K4me3. Initially we examined the bulk levels of H3K4me3 by western blot after SET1 protein depletion (Fig.2a and Extended Data Fig.2a). Surprisingly, despite widespread effects on gene expression, we observed only a very modest and non-significant reduction in H3K4me3, even after several days of SET1 protein depletion^33^. However, our bulk western analysis may simply overlook gene-specific effects on H3K4me3. Therefore, we also carried out cChIP-seq for H3K4me3 at 2, 4 and 24 hours after SET1 protein depletion (Fig.2b-c and Extended Data Fig.2b-d). Again, this revealed only very modest reductions in H3K4me3 at gene promoters and there was no significant correlation between the effects on H3K4me3 and reductions in gene expression (Fig.2b and d and Extended Data Fig.2e). This argues that the SET1 complexes are not responsible for placing the majority of H3K4me3 at gene promoters in ESCs, and suggests that the effects we observe on gene expression after depletion of SET1 complexes may be independent of H3K4me3 and SET1 complex methyltransferase activity.

Based on these findings, we sought to more directly examine whether SET1 complexes regulate gene expression independently of their methyltransferase activity. To achieve this, we developed a chromatinised gene reporter system containing a single copy transgene in which Tet operator DNA binding sites (TetO) were coupled to a minimal gene promoter that drives luciferase expression (Fig.2e). Fusing a protein of interest to the reverse Tet Repressor DNA binding domain (rTetR) enables its recruitment to the TetO array upon the addition of doxycycline (Fig.2e). Consistent with a role for SET1 complexes in supporting gene expression, tethering wild type SET1A to the promoter resulted in increased reporter gene expression (Fig.2f). Interestingly, tethering a version of SET1A in which we had mutated key residues required for its methyltransferase activity supported gene expression in a manner that was similar to the wild type protein (Fig.2f and Extended Data Fig.2f-g). Together, our histone modification analysis and tethering experiments suggest that alterations in gene expression observed after SET1 protein depletion do not primarily manifest from a loss of H3K4me3 and that SET1A can support gene expression independently of its methyltransferase activity.

### SET1 complexes support gene expression through an interaction with WDR82

Given that SET1A can support gene expression independently of its methyltransferase activity, we set out to determine what region of the protein was responsible for this effect. To address this, we took advantage of our reporter gene system and tested the capacity of various SET1A fragments to support gene expression. These fragments included: the conserved N-terminal region that has been proposed to interact with WDR82^18, 82^; the RRM domain which, in other proteins, can interact with RNA^83^; the central region of SET1A which lacks significant sequence conservation; and the N-SET/SET domain which interacts with CFP1/WDR5/RBBP5/ASH2L/DPY30 and is required for chromatin binding and methyltransferase activity (Fig.3a)^17^. Interestingly, only the short N-terminal domain (NTD) of SET1A was sufficient to support gene expression, and it did so to a similar extent as the full-length protein (Fig.3b and Extended Data Fig.3b). This suggests that the other domains in SET1A, and their interacting proteins, do not contribute significantly to supporting gene expression in this context. Furthermore, the equivalent NTD of SET1B was also sufficient to support gene expression, indicating that this activity is conserved amongst SET1 paralogues (Extended Data Fig.3a-b).

Having shown that the NTDs of SET1 proteins are sufficient to support gene expression, we hypothesised that this effect may rely on their capacity to interact with WDR82. To test this possibility, we carried out sequence conservation analysis of the NTD across SET1 orthologues to identify a putative WDR82 interaction motif (Fig.3c). This revealed a highly conserved trio of amino acids, herein referred to as the DPR motif. When the DPR motif was mutated to alanines (DPR/AAA), the SET1A NTD was unable to interact with WDR82 (Fig.3d). Importantly, when we tethered the DPR/AAA-NTD of SET1A to the reporter gene promoter, it was unable to support gene expression (Fig.3e). Therefore, a highly conserved DPR motif in the N-terminal domains of SET1 proteins interacts with WDR82, and this interaction can support gene expression.

### SET1 complexes support transcription over CpG islands

Having discovered that SET1A can regulate gene expression through binding to WDR82, and knowing that WDR82 can interact with RNA Pol II^18^, we reasoned that SET1 complexes may directly influence RNA Pol II function and gene transcription. To explore this possibility, we examined transcription using calibrated transient transcriptome sequencing (cTT-seq) and RNA Pol II occupancy using cChIP-seq (Fig.4a).

In agreement with cRNA-seq analysis, we observed a widespread reduction in cTT-seq signal after depletion of SET1 proteins, with 3098 genes, or ∼25% of all transcribed genes, displaying a significant reduction in transcription (Fig.4b and Extended Data Fig.4a-b). In contrast, only 75 genes exhibited a significant increase in transcription. Again, in line with our findings from gene expression analysis using cRNA-seq, SET1-dependent genes were also more moderately transcribed compared to SET1-independent genes (Extended Data Fig.4c). This analysis demonstrates that the SET1 complexes almost exclusively support gene transcription.

To better understand how SET1 complexes might influence the function of RNA Pol II, we next examined RNA Pol II occupancy by cChIP-seq. Interestingly, at both SET1-dependent and SET1-independent genes, we observed only very modest effects on RNA Pol II binding over the region corresponding to the transcription start site and promoter proximal pause site (Fig.4c-d and Extended Data Fig.4d). This suggests that SET1 complexes do not play a central role in influencing RNA Pol II occupancy during the very earliest stages of transcription. However, we did observe reductions in RNA Pol II occupancy in the body of SET1-dependent genes, an effect that was not evident at SET1-independent genes (Fig.4c-d and Extended Data Fig.4d). This suggests that SET1 complexes may support RNA Pol II in transcribing into the gene body.

To investigate this possibility in more detail, we examined our cTT-seq data, which also provides spatial information about the level of transcription across genes^84, 85^. Interestingly, when SET1 proteins were depleted, we discovered that SET1-dependent genes still initiated transcription from the gene promoter, albeit at reduced levels (Fig.4e-f and Extended Data Fig.4e). However, transcription then rapidly attenuated within the CGI in a region ∼500-1500 bp downstream of the TSS, and this tended to be skewed towards the 3’ boundary of the CGI (Fig.4f-g). Given that we did not observe accumulation of RNA Pol II in the promoter region or the body of SET1-dependent genes in RNA Pol II cChIP-seq after SET1 protein depletion, the reduction in transcription does not appear to be due to increased promoter proximal pausing or reduced elongation rate in the gene body (Fig.4d)^86^. Therefore, we propose that the observed attenuation of transcription is likely caused by premature transcription termination (PTT), which would be consistent with largely unaffected promoter-proximal occupancy of RNA Pol II and a reduction of RNA Pol II in the body of SET1-dependent genes. PTT might also in part explain the reduced cTT-seq signal observed at the 5’ end of SET1-dependent genes following SET1 protein depletion, since the products of PTT are often rapidly degraded by the exosome^5, 58^. Together these observations suggest that SET1 complexes bind to CGIs, which tend to stretch in to the 5’ end of genes, where they counteract the attenuation of early transcription. Furthermore, this activity appears to be particularly important for the expression of moderately transcribed genes.

### SET1 complexes counteract premature transcription termination by ZC3H4

Given the attenuation of transcription over CGIs after depletion of SET1 proteins, we reasoned that SET1 complexes may enable transcription by counteracting the activity of an opposing PTT pathway. We have shown that SET1 complexes can support gene expression via an interaction with the RNA Pol II-binding protein, WDR82. Interestingly, WDR82 also interacts with ZC3H4, which has been shown to localise with RNA Pol II and contribute to transcription termination, particularly of extragenic transcription^13, 45, 68-70^. Therefore, we hypothesised that the enrichment of WDR82-containing SET1 complexes at CGIs may support transcription of moderately transcribed genes by counteracting the termination of early transcription by WDR82-containing ZC3H4 complexes.

To explore this possibility, we first epitope-tagged the endogenous ZC3H4 gene and carried out ChIP-seq analysis to examine binding of ZC3H4 in the genome (Fig.5a-b and Extended Data Fig.5a). As reported previously, we found that ZC3H4 localises with RNA Pol II at actively transcribed genes and at regions of extragenic transcription^68, 69^ (Fig.5a-b and Extended Data Fig.5b). However, when we examined ZC3H4 binding in more detail a number of important features emerged. Firstly, despite its capacity to interact with RNA Pol II, ZC3H4 does not localise precisely to the peak of RNA Pol II at transcription start sites, but is instead enriched on the shoulders of RNA Pol II peaks (Fig.5a-b). Secondly, at SET1-independent genes, ZC3H4 is enriched upstream of gene promoters, coincident with the location of antisense transcripts and consistent with a reported role for ZC3H4 in terminating antisense extragenic transcription (Fig.5a-b). Finally, at SET1-dependent genes, we observed a distinct enrichment of ZC3H4 in coding regions, just downstream of the transcription start site (Fig.5a-b). The location of this enrichment is coincident with the observed attenuation of transcription after the depletion of the SET1 proteins, and with the underlying CGI element (Fig.4g). Therefore, despite previous reports that ZC3H4 primarily affects extragenic and non-coding RNAs, we hypothesised that ZC3H4 might also play a prominent role in PTT at protein-coding genes.

To test this possibility, we engineered a dTAG into the endogenous ZC3H4 gene. As was the case for SET1A/B, dTAG13 treatment resulted in a near complete depletion of ZC3H4 within 2 hours (Fig.5c). To examine whether ZC3H4 depletion affects transcription, we carried out cTT-seq. In the absence of ZC3H4, extragenic upstream antisense transcription from promoters was increased, as was enhancer transcription, consistent with its proposed role in terminating these transcripts in other cell types (Fig.5d)^68, 69^. Importantly, however, we also observed that the depletion of ZC3H4 resulted in the increased transcription of 2599 genes, indicating that ZC3H4 also significantly influences genic transcription (Fig.5e). Interestingly, SET1-dependent genes displayed an increase in transcription in the absence of ZC3H4, whereas SET1-independent genes tended to show a slight reduction in transcription (Fig.5f-g and Extended Data Fig.5c). This suggests that ZC3H4 can attenuate transcription of protein-coding genes, and that this effect is most obvious at SET1-dependent genes. Importantly, this indicates that SET1 and ZC3H4 complexes have seemingly opposing effects on transcription at SET1-dependent genes.

Based on these findings, we wanted to test whether ZC3H4 complexes were responsible for the profound attenuation of gene transcription observed when SET1 proteins were depleted. To address this important question, we engineered a dTAG into the ZC3H4 gene in the dTAG-SET1A/B cell line (Extended Data Fig.5d). We then treated these cells with dTAG13 to simultaneously deplete both SET1 and ZC3H4 complexes and carried out cTT-seq. Remarkably, this revealed that the attenuation of gene transcription caused by depletion of SET1 proteins was completely reversed when ZC3H4 was also depleted (Fig.5h-k and Extended Data Fig.5e). In contrast, antisense transcription upstream of genes and enhancer-associated transcription, both of which infrequently overlap with CGIs and show little occupancy of SET1 complexes (Extended Data Fig.5b), were highly susceptible to attenuation by ZC3H4 (Extended Data Fig.5f-g). Therefore, we propose that ZC3H4 complexes have widespread roles in causing PTT both of genic and extragenic transcription. However, CGIs, which are associated with the 5’ ends of genes and nucleate SET1 complexes, preferentially protect genic transcription from PTT as a means to enable gene expression.

## Discussion

How CGIs regulate transcription has remained an elusive and enigmatic component of vertebrate gene regulation. Furthermore, while PTT has recently emerged as an important regulator of gene transcription, the mechanisms through which it is controlled remain poorly understood and represent a major conceptual gap in our understanding of gene regulation. Here we discover that CGI-associated SET1 complexes primarily function to enable gene expression (Fig.1). Unexpectedly, this is not mediated by their H3K4me3 methyltransferase activity (Fig.2), but instead relies on their capacity to interact with the RNA Pol II-binding protein WDR82 (Fig.3). Removal of SET1 complexes causes moderately transcribed genes to become sensitive to PTT within a zone downstream of the TSS that is coincident with the underlying CGI (Fig.4). We discover that this PTT is driven by ZC3H4 complexes, which also contain WDR82. This suggests that SET1 complexes may function at CGIs through WDR82 to counteract PTT by ZC3H4 complexes (Fig.5). In agreement with this, simultaneous removal of SET1 and ZC3H4 complexes reverses the requirement for SET1 complexes in gene transcription, demonstrating that SET1 complexes primarily function through binding to CGIs to counteract ZC3H4-dependent PTT (Fig.5). Therefore, we uncover an unexpected new gene regulatory mechanism whereby a CGI-binding complex at actively transcribed genes acts to counteract PTT and enable gene expression.

**Fig.2:**
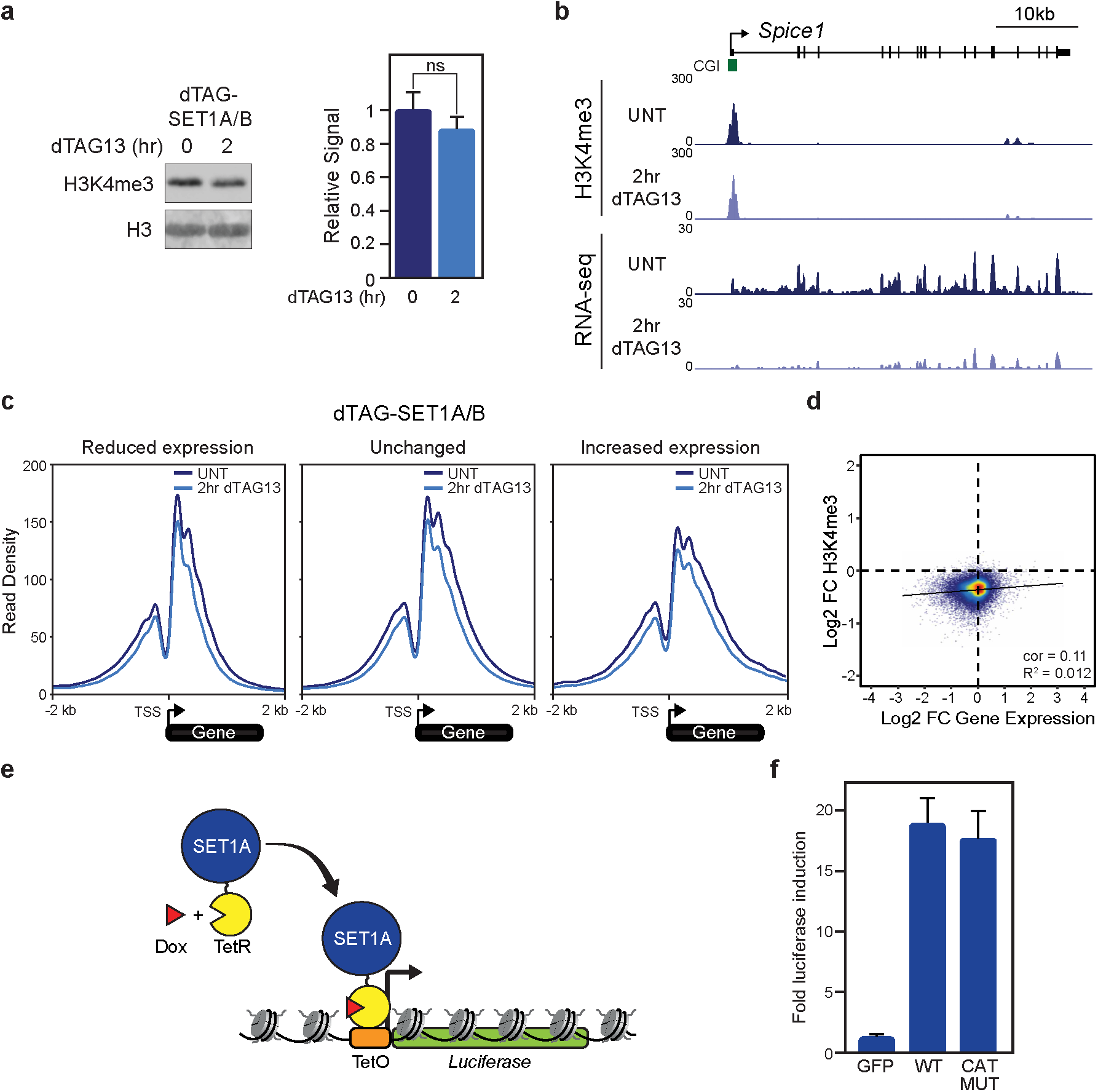
SET1 complexes can regulate gene expression independently of H3K4me3 and methyltransferase activity. **(a)** Western blot analysis of H3K4me3 levels in the untreated dTAG-SET1A/B line and following 2 hours of dTAG13 treatment (left panel). H3 is included as a loading control. Quantitation of the alteration in H3K4me3 levels after dTAG13 treatment from 2 biological replicates. The error bars represent standard error of the mean (SEM) (right panel). ns indicates that the quantified change is not significant. **(b)** A genomic snapshot comparing H3K4me3 cChIP-seq signal (top panels) and cRNA-seq (bottom panels) before and after 2 hours of dTAG13 treatment in the dTAG-SET1A/B line at the *Spice1* gene. **(c)** Metaplot analysis of H3K4me3 cChIP-seq around the transcription start site (TSS) of genes the have reduced expression (left panel), unchanged expression (middle panel), or increased expression (right panel) in the dTAG-SET1A/B line before (dark blue lines) and after 2 hours of dTAG13 treatment (light blue lines). Only expressed genes are included (reduced expression, n=2544; unchanged, n=9989; increased expression, n=495). **(d)** A scatter plot comparing the log2 fold change (log2FC) in H3K4me3 cChIP-seq signal and cRNA-seq signal in the dTAG-SET1A/B line following 2 hours of dTAG13 treatment. The correlation (cor) and R^2^ values are indicated. Only genes that have a peak of H3K4me3 in untreated cells are included (n=14065). **(e)** A schematic illustrating the chromatinised reporter gene. TetO binding sites are coupled to a minimal core promoter and a luciferase reporter gene. rTetR-fusion proteins are tethered to the reporter gene by the addition of doxycycline (Dox) and effects on gene expression can be monitored. **(f)** A bar plot showing the fold induction of reporter gene expression following tethering of GFP, SET1A (WT) and SET1A with catalytic mutations in its SET domain (MUT). Error bars represent SEM from three biological replicates.

**Fig.3:**
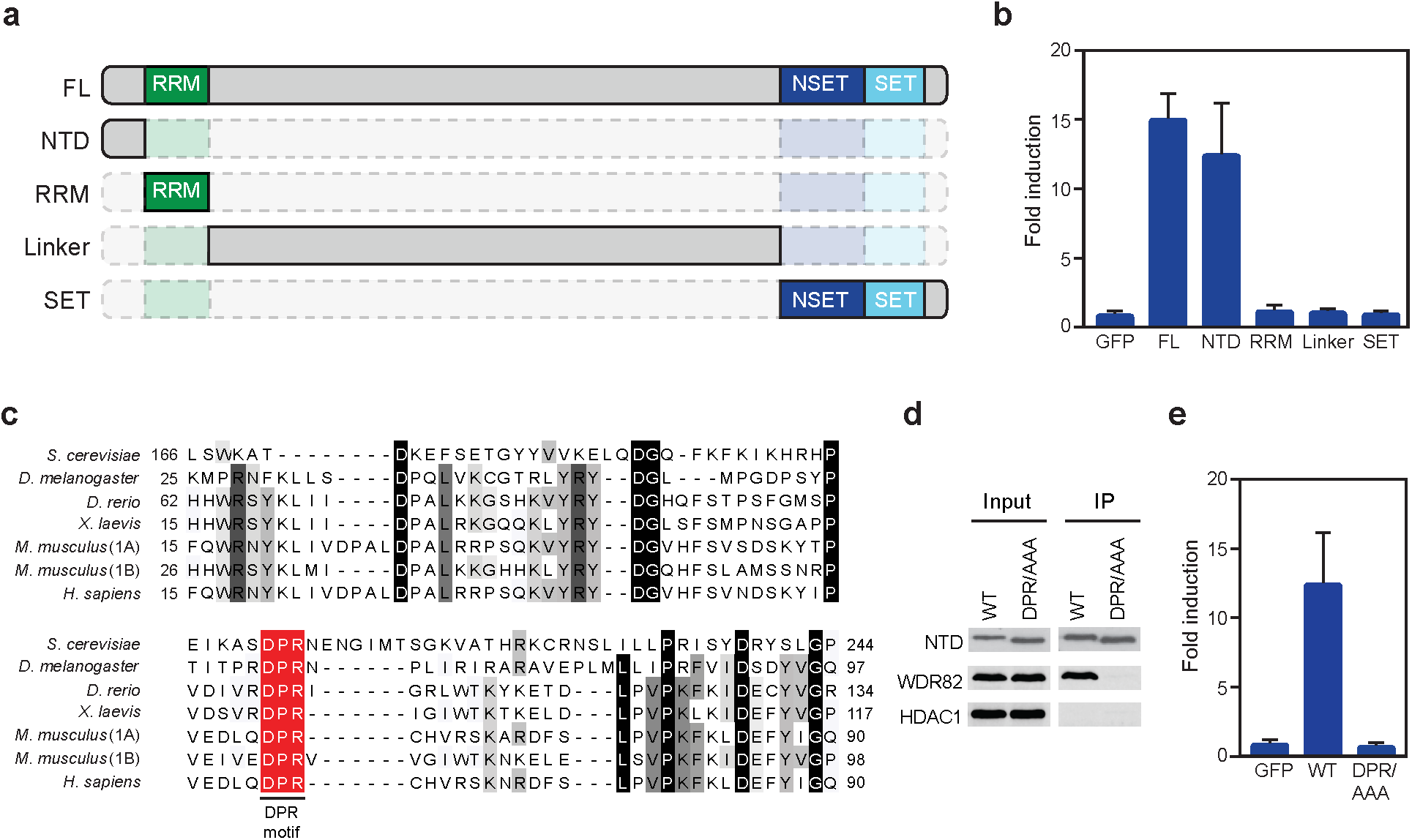
SET1 complexes support gene expression through an interaction with WDR82. **(a)** A schematic illustrating the SET1A domains that were tethered to the reporter gene. **(b)** A bar plot showing the fold induction of reporter gene expression following tethering of GFP, full length SET1A (FL), NTD, RRM, Linker, and the NSET-SET (SET) domain fragments to the reporter gene. Error bars represent SEM from five biological replicates. **(c)** A multiple sequence alignment of the N-terminal domains of SET1A (1A) or SET1B (1B) from the indicated species. The red box highlights the conserved and invariant DPR motif. **(d)** An immunoprecipitation (IP) of the NTD of WT SET1A or the DPR/AAA mutant followed by western blot for WDR82. HDAC1 functions as a loading control for the Input samples and a negative control for interaction with SET1A. **(e)** A bar plot showing the fold induction of reporter gene expression following tethering of GFP, WT-NTD, or the DPR/AAA-NTD of SET1A. Error bars represent SEM from five biological replicates.

**Fig.4:**
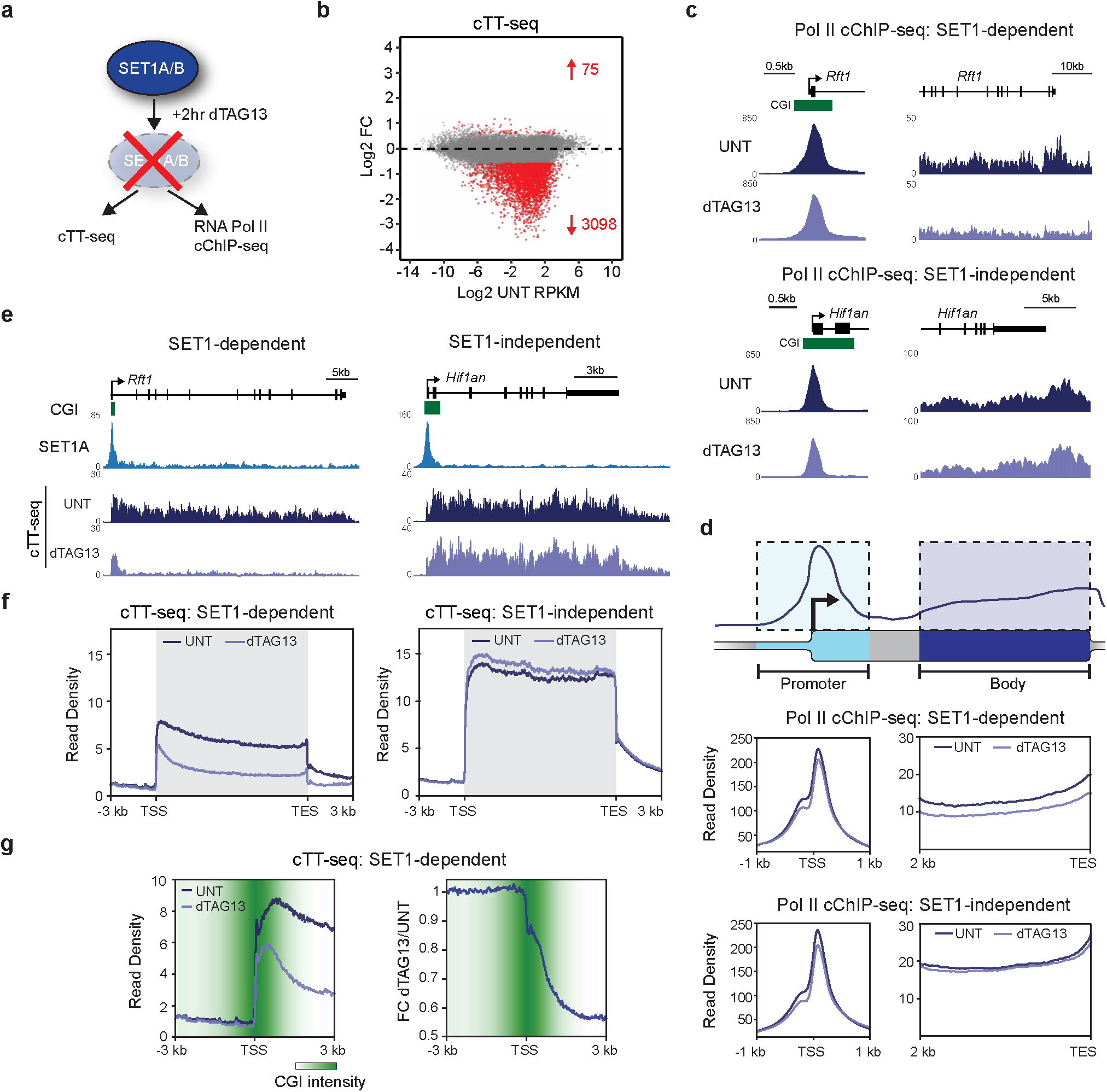
SET1 complexes support transcription over CpG islands. **(a)** A schematic illustrating the experiments carried out in the dTAG-SET1A/B line to examine RNA Pol II occupancy (Pol II cChIP-seq) and transcription (cTT-seq). **(b)** An MA-plot showing log2 fold changes in transcription (cTT-seq) in the dTAG-SET1A/B line following 2hr of dTAG13 treatment (n=20633). Significant changes in transcription (p-adj<0.05 and >1.5-fold) are coloured red and the number of significantly changed genes is indicated. **(c)** Genomic snapshots of RNA Pol II occupancy (Pol II cChIP-seq) at a SET1-dependent (upper panel, *Rft1*) and SET1-independent (lower panel, *Hif1an*) gene in untreated (UNT) cells (dark purple) or cells treated with dTAG13 for 2 hours (light purple). The left hand panels correspond to gene promoter occupancy and the right hand panels to gene body occupancy. **(d)** The top panel is a cartoon schematic illustrating the typical RNA Pol II cChIP-seq signal over a gene with the gene promoter region highlighted in light blue and the gene body in light purple. The bottom panels correspond to metaplot analysis of RNA Pol II cChIP-seq signal in the dTAG-SET1A/B line in cells that are either untreated (UNT) or treated with dTAG13 for 2 hours. The RNA Pol II cChIP-seq signal corresponding to the gene promoter and body regions (see schematic in top panel) of all transcribed SET1-dependent genes (middle panel, n=2633) and SET1-independent genes (bottom panel, n=9151) is shown. **(e)** Genomic snapshots of cTT-seq signal in the dTAG-SET1A/B line at a SET1-dependent (left panel, *Rft1*) and SET1-independent (right panel, *Hif1an*) gene in untreated (UNT) cells (dark purple) or cells treated with dTAG13 for 2 hours (light purple). The location of CGIs is shown in green and SET1A cChIP-seq signal in light blue. **(f)** Metaplot analysis of transcription (cTT-seq) in the dTAG-SET1A/B line in cells that are either untreated (UNT, dark purple) or treated with dTAG13 for 2 hours (light purple) for all actively transcribed SET1-dependent genes (left panel, n=2633) and SET1-independent genes (right panel, n=9151). **(g)** Metaplot analysis of transcription (cTT-seq) in the dTAG-SET1A/B line zoomed in to the transcription start site (TSS) of all actively transcribed SET1-dependent genes (left panel, n=2633). Also shown is a metaplot analysis of the fold change in cTT-seq signal between dTAG13-treated and untreated dTAG-SET1A/B cells at the transcription start site (TSS) of all SET1-dependent genes (right panel, n=2633). For both plots, the log2-transformed average intensity of non-methylated DNA is indicated as a proxy for the location of the CGI.

**Fig.5:**
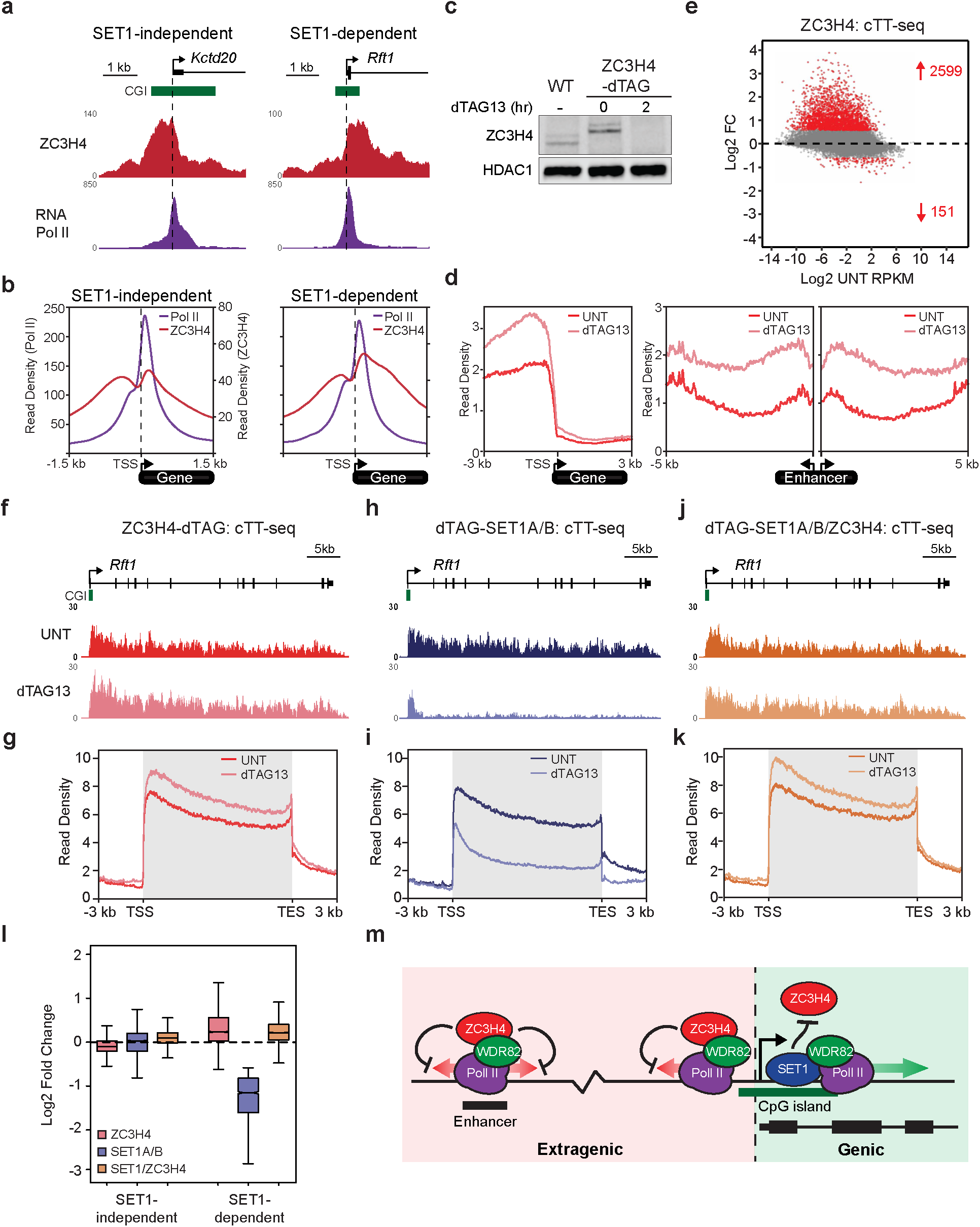
SET1 complexes counteract premature transcription termination by ZC3H4. **(a)** Genomic snapshots of ZC3H4 and RNA Pol II ChIP-seq signal at a SET1-independent (*Kctd20*) and SET1-dependent (*Rft1*) gene. The location of CGIs are indicted by green boxes. **(b)** Metaplot analysis of ZC3H4 and RNA Pol II ChIP-seq at the TSS of transcribed SET1-independent (n=9151) and SET1-dependent genes (n=2633). The read density of RNA Pol II is shown on the left axis and read density of ZC3H4 on the right axis. **(c)** A western blot comparing ZC3H4 levels in wild type (WT) cells and the ZC3H4-dTAG line. For the ZC3H4-dTAG line treatment with dTAG13 for 2 hours leads to depletion of ZC3H4. HDAC1 functions as a loading control. **(d)** Metaplot analysis of transcription (cTT-seq) in the ZC3H4-dTAG line that is either untreated (dark red) or treated with dTAG13 for 2 hours (light red), showing upstream antisense transcription at all TSSs (left panel, n=20633) and enhancer transcription (right panels, n=4156). **(e)** An MA-plot showing log2 fold changes in transcription (cTT-seq) in the ZC3H4-dTAG line following 2hr treatment with dTAG13 (n=20633). Significant changes in transcription (p-adj<0.05 and >1.5-fold) are coloured red and the number of significantly changed genes is indicated. **(f)** A genomic snapshot of cTT-seq signal in the ZC3H4-dTAG line at a SET1-dependent (*Rft1*) gene in untreated cells (UNT) or cells treated with dTAG13 for 2 hours. **(g)** Metaplot analysis of transcription (cTT-seq) in the ZC3H4-dTAG line in cells that are either untreated (UNT) or treated with dTAG13 for 2 hours at all transcribed SET1-dependent genes (n=2633). **(h-i)** As per (f) and (g) but for dTAG-SET1A/B cells. **(j-k)** As per (f) and (g) but for dTAG-SET1A/B/ZC3H4 lines. **(l)** A box plot showing the log2 fold change in transcription at transcribed SET1-independent (n=9151) and SET1-dependent (n=2633) genes for the dTAG-SET1A/B, dTAG-ZC3H4, and dTAG-SET1A/B/ZC3H4 lines after treatment with dTAG13 for 2 hours. **(m)** A cartoon illustrating a model whereby WDR82-containing SET1 complexes bind to CGIs to enable genic transcription by counteracting premature transcription termination by WDR82-containing ZC3H4 complexes. In contrast, extragenic transcription that emanates from regions lacking CGIs and SET1 complex occupancy is subject to termination by WDR82-containing ZC3H4 complexes. In this model, CGIs and SET1 complex occupancy would distinguish genic from extragenic transcription, and protect genic transcription from premature transcription termination to enable gene expression.

In contextualising these new discoveries, it is important to consider how ZC3H4 and SET1 complexes interface with their target genes and hence the logic that might underpin how their functional integration ensures appropriate gene expression. Key to this is likely the fact that ZC3H4 and SET1 complexes have a shared interaction partner, WDR82^13, 45^. WDR82 preferentially binds the C-terminal heptapeptide repeat (CTD) of the largest subunit of RNA Pol II when it is phosphorylated on serine 5 (Ser5P), which occurs when initiated RNA Pol II transitions into elongation^18, 44^. Although we currently have a limited understanding of the precise mechanisms that enable ZC3H4 complex targeting, in addition to binding CTD-Ser5P via WDR82, ZC3H4 can also interact with the nuclear RNA cap-binding complex protein ARS2^87^. Based on these interactions, we propose that ZC3H4 complexes may generically recognise early elongating Pol II through binding to CTD-Ser5P and an exposed and capped RNA, and then function to attenuate transcription. In agreement with this possibility, we identify ZC3H4 binding primarily in regions flanking TSSs where Pol II has initiated (Fig.5a-b), and detect a widespread influence on both genic and extragenic transcription when it is depleted.

If ZC3H4 complexes provide a generic mechanism to attenuate early transcription, this would be beneficial for limiting the production of extragenic transcripts from enhancers and upstream of gene promoters, which, in many cases, are non-functional^68, 69^. However, genic transcription would also be susceptible to the influence of ZC3H4 complexes and this could be highly detrimental to gene expression. Importantly, we now show that SET1 complex binding to CGIs of actively transcribed genes counteracts PTT by ZC3H4 complexes. Like ZC3H4 complexes, SET1 complexes interact with WDR82, and this interaction appears to be important for their effects on gene expression. We therefore speculate that WDR82-containing SET1 complexes may counteract the activity of ZC3H4 at actively transcribed CGI-associated genes by competing for binding to early elongating RNA Pol II and thus limiting the capacity of ZC3H4 to cause PTT. As such, SET1 complex occupancy at gene promoter-associated CGIs would help to distinguish genic transcription from extragenic transcription, which is not typically associated with CGIs, thereby limiting PTT to the latter^7, 88^ (Fig.5m). Given that CGIs tend to stretch into the 5’ end of genes, this could explain why we find that sense genic transcription is preferentially protected from PTT by SET1 complexes, while antisense transcription is susceptible. Such a mechanism could also explain previous findings that RNA Pol II is less susceptible to PTT within CGI elements, whilst high-throughput screens identified CGI DNA as a potent inhibitor of PTT^54, 56^. However, an outstanding question remains as to why the anti-termination activity of SET1 complexes appears to be particularly important for the transcription of moderately transcribed genes. One possible explanation may be that additional mechanisms involved in driving high levels of gene transcription impart additional transcription processivity influences that also limit PTT by ZC3H4 complexes, a possibility that will be important to test in the context of future studies.

In conclusion, we identify a new gene regulatory mechanism whereby SET1 complexes bind to actively transcribed CGIs to specifically counteract PTT by ZC3H4 complexes. Furthermore, we demonstrate that termination of early transcription by ZC3H4 complexes is widespread, and show that pervasive PTT must be counteracted to support normal gene expression. Importantly, this also reveals a new role for CGIs in distinguishing genic from extragenic transcription, and protecting genes from premature transcription termination to enable normal gene expression.

## Acknowledgments

We would like to thank Tom Milne, Krzysztof Kus, Lars Jansen, and Alan Au for critical reading of the manuscript, as well as members of the Klose lab for fruitful discussions. We thank Angelika Feldmann, Nadezda Fursova, and Paula Dobrinic for helpful discussions regarding computational analysis. We are grateful to Amanda Williams at the Department of Zoology, Oxford, for sequencing support on the NextSeq 500. We would like to thank Behnam Nabet, Nathanael Gray, Carole Bataille and Angela Russell for generous provision of dTAG13 compound. We thank Martin Houlard for providing the T7-Scc1 cell line and Michael Rehli for providing the pCpGL-CMV/EF1 plasmid. Work in the Klose lab is supported by the Wellcome Trust (209400/Z/17/Z). A.L.H (203829/Z/16/A) and A.H.T (102349/Z/13/Z) were supported by a Wellcome Trust studentships. J.R.K is supported by the Oxford-Wolfson Marriott Graduate Scholarship.

## Author Contributions

A.L.H and R.J.K conceived the project and wrote the manuscript with contributions from all co-authors. A.L.H performed most of the experiments, data analysis, and visualisation. A.T.S performed the smRNA-FISH, data analysis, and visualisation. J.R.K performed the WDR82 interaction analysis, multiple sequence analysis, and visualisation. A.H.T generated the gene expression reporter cell line. A.L, E.D, N.P.B contributed to cell line generation, experimentation, and drafting the manuscript. N.P.B contributed to visualisation of data in figures. R.J.K supervised the project.

## Materials and Methods

### Cell culture

Mouse embryonic stem cells were grown in Dulbecco’s Modified Eagle Medium (Thermo Fisher Scientific) supplemented with fetal bovine serum (FBS, 15% Biosera or 10% Sigma), 1x non-essential amino acids (Thermo Fisher Scientific), 2 mM L-glutamine (Thermo Fisher Scientific), 1x penicillin/streptomycin (Thermo Fisher Scientific), 0.5 mM beta-mercaptoethanol (Thermo Fisher Scientific), and 10 ng/ml leukaemia inhibitory factor (produced in-house). ESCs were grown on gelatinised plates at 37⁰C and 5% CO2. Cell lines expressing dTAG fusion proteins were treated with 100 nM dTAG-13 (produced by Behnam Nabet and Nathanael Gray^76^ or Carole Bataille and Angela Russell) to induce protein depletion.

*Drosophila melanogaster* SG4 cells were grown adhesively at 25°C in Schneider’s *Drosophila* Medium (Thermo Fisher Scientific) supplemented with 1x penicillin/streptomycin and 10% heat-inactivated FBS (Biosera). Human HEK 293T cells were grown at 37⁰C and 5% CO2 in Dulbecco’s Modified Eagle Medium, supplemented with 10% FBS (Biosera), 1x penicillin/streptomycin, 2 mM L-glutamine and 0.5 mM beta-mercaptoethanol. All cell lines were routinely tested to ensure they were mycoplasma free.

### Stable cell line generation

To allow for their rapid depletion, we introduced an N-terminal dTAG into the endogenous SET1A and SET1B genes, and a C-terminal dTAG into the endogenous ZC3H4 gene. To allow for efficient chromatin immunoprecipitation, we also introduced an N-terminal 3xT7-2xStrepII-dTAG tag into the endogenous SET1A gene. To generate the luciferase reporter cell line, we modified a previously described mouse ESC line containing a single copy insertion of a human gene desert bacterial artificial chromosome^89^ with a cassette containing 7 CpG-free TetO DNA binding sites^89^, followed by a CpG-free Ef1a promoter and a luciferase reporter gene^90^.

Stable cell lines were engineered using CRISPR/Cas9-mediated genome editing. sgRNAs were designed using the CRISPOR online tool (http://crispor.tefor.net) and oligonucleotides encoding sgRNAs were cloned into the pSpCas9(BB)-2A-Puro plasmid as previously described (Addgene #62988)^91^. Targeting constructs containing the sequence to be inserted and approximately 500 bp homology arms were cloned by Gibson Assembly (NEB). The FKBP12^F36V^ tag (dTAG) was obtained from Addgene (#91797). ESCs were transfected at approximately 70% confluency in a 6-well plate with 0.5 µg of guide plasmid and 2 µg of targeting construct using Lipofectamine 3000, according to the manufacturer’s protocol (Thermo Fisher Scientific). The following morning, transfected cells were passaged to new plates at low density and selected with 1 ug/ml puromycin for 48 hr. Individual colonies were picked into 96-well plates and positive clones identified by PCR screens of genomic DNA.

### Luciferase reporter assays

To express rTetR-fusion proteins, fragments comprising GFP, full length SET1A and the SET1A catalytic mutant were Gibson assembled into a plasmid backbone containing a CAG promoter, N-terminal FLAG-StrepII tag, nuclear localisation signal and rTetR, as described previously^89^. The rTetR was replaced with that from TetON-3G plasmid for all SET1A minimal domain fusions and full length SET1A and SET1B NTD in Fig. 3 (Addgene plasmid #96963)^92, 93^.

ESCs were transfected at approximately 70% confluency with Lipofectamine 2000, according to the manufacturer’s protocol (Thermo Fisher Scientific). For preparation of nuclear extract, cells were transfected in 6-well plates with 2.5 µg plasmid. For luciferase assays, cells were transfected in 96-well plates with 98 ng rTetR plasmid and 2 ng pRL *Renilla* Luciferase control reporter plasmid (Promega) to control for transfection efficiency. Each transfection was performed in six wells of a 96-well plate to obtain three technical replicates for both untreated and treated with doxycycline. Following overnight transfection of plasmids expressing rTetR-fusion proteins and *Renilla* luciferase, three wells for each transfection were treated with 1 µg/ml doxycycline for 6 hr. Luciferase reporter assays were performed with the Dual-Luciferase Reporter Assay System (Promega). In brief, cells were lysed in 20 µl 1x passive lysis buffer by shaking at room temperature for 20 min. 10 µl cell lysate was added to 50 µl Luciferase Assay Reagent II and Firefly luciferase measured using a 10 second measurement in a Luminometer. 50 µl Stop & Glo Reagent was added and *Renilla* luciferase measured using a 10 second measurement. Each Firefly reading was normalised to the respective *Renilla* reading. Technical replicates were averaged and normalised to the readings obtained in the absence of doxycycline. Each assay was performed in at least biological triplicates.

### Protein extraction and immunoblotting

To prepare histone extracts, pelleted cells were washed in RSB (10 mM Tris HCl pH 7.4, 10 mM NaCl, 3 mM MgCl2), centrifuged at 240xg for 5 min and resuspended in RSB supplemented with 0.5% NP-40. Following incubation on ice for 10 min, cells were centrifuged at 500xg for 5 minutes. The nuclear pellet was resuspended in 5 mM MgCl2, an equal volume of 0.8 M HCl added, and then incubated on ice for 20 min to extract histones. After centrifugation for 20 min at 18,000xg, the supernatant was taken and histones precipitated by adding TCA to 25% v/v and incubating on ice for 30 min. Histones were pelleted by centrifugation at 18,000xg for 15 min, and the pellet was washed twice in cold acetone. The histone pellet was resuspended by gentle vortexing in 1x SDS loading buffer (2% SDS, 100 mM Tris pH 6.8, 100 mM DTT, 10% glycerol, 0.1% bromophenol blue) and boiling at 95⁰C for 5 min. Any insoluble precipitate was pelleted by centrifugation at 18,000xg for 15 min and the soluble fraction taken as the histone extract. Histone extract concentrations were compared across samples by SDS-polyacrylamide gel electrophoresis (SDS-PAGE) and Coomassie Blue staining.

To prepare nuclear extracts, cell pellets were resuspended in 10x pellet volumes of buffer A (10 mM HEPES pH 7.9, 1.5 mM MgCl2, 10 mM KCl, 0.5 mM DTT, 0.5 mM PMSF, 1x cOmplete protease inhibitor cocktail (PIC, Roche)) and incubated for 10 min on ice. After centrifugation at 500xg for 5 min, the cell pellet was resuspended in 3x pellet volumes of buffer A supplemented with 0.1% NP-40 and incubated on ice for 10 min. Nuclei were pelleted at 1500xg for 5 min and then resuspended in 1x pellet volume of buffer C (250 mM NaCl, 5 mM HEPES pH 7.9, 26% glycerol, 1.5 mM MgCl2, 0.2 mM EDTA, 0.5 mM DTT, 1x PIC). The volume of the nuclear suspension was measured and the NaCl concentration increased to 400 mM by dropwise addition of 5 M NaCl. Nuclei were incubated at 4⁰C for 1 hr with gentle inversion to extract nuclear proteins. After centrifugation at 18,000xg for 20 min, nuclear proteins were recovered in the supernatant. Protein concentration was determined by Bradford assay (BioRad) and typically 25 µg was used for western blotting.

Protein extracts were resolved using either home-made SDS-PAGE gels or 3-8% NuPAGE Tris-Acetate gels (Thermo Fisher Scientific), when analysing proteins of a molecular weight >180 kDa. Typically, proteins were transferred to nitrocellulose membrane by semi-dry transfer using the Trans-Blot Turbo Transfer System (BioRad). Transfer was performed as per the manufacturer’s guidelines, depending on the size of the proteins being transferred. Membranes were imaged using an Odyssey Fc system (LI-COR). Changes in bulk protein levels were quantified relative to those of loading controls. For SET1B western blots, proteins were transferred to 0.45 µm nitrocellulose membrane by wet transfer. Transfer was performed in 1x wet transfer buffer (25 mM Tris, 192 mM glycine, 10% Methanol, 1% SDS) at 100 V for 90 min at 4⁰C. Membranes were developed by chemiluminescence.

Antibodies used for western blot analysis were anti-SET1A (Bethyl Laboratories, A300-289A), anti-SET1B (Cell signalling, D1U5D), anti-SUZ12 (Cell Signalling, D39F6), anti-WDR5 (Cell Signalling, D9E1I), anti-RBBP5 (Cell Signalling, D3I6P), anti-ASH2L (Cell Signalling, D93F6), anti-WDR82 (Cell Signalling D2I3B), anti-CFP1 (Klose Lab), anti-H3K4me3 (Klose Lab), anti-H3 (Cell Signalling, 96C10), anti-FS2 (Klose Lab), anti-RING1B (Klose Lab), anti-FLAG (Sigma, A8592), anti-HDAC1 (Abcam, ab109411), anti-ZC3H4 (Atlas Antibodies, HPA040934).

### Co-Immunoprecipitation

Immunoprecipitations (IPs) of SET1A were performed using 550 µg of nuclear extract. Extracts were diluted to 500 µl with BC150 (150 mM KCl, 10% glycerol, 50 mM HEPES pH 7.9, 0.5 mM EDTA, 0.5 mM DTT and 1x PIC), with 250 units Benzonase Nuclease (Sigma). Protein A beads (Repligen) were blocked in BC150 supplemented with 1% Fish gelatin (Sigma) and 0.2 mg/ml BSA (NEB) for 1 hr at 4⁰C. Extracts were pre-cleared with 50 µl slurry blocked beads for 1 hr at 4⁰C and then incubated rotating with 20 µl SET1A antibody (Klose Lab) overnight at 4⁰C. 50 µl slurry blocked Protein A beads were used to precipitate antibody-bound protein at 4⁰C for 3 hr. Beads were pelleted at 1000xg, and washed 3 times with BC150 + 0.02% NP-40, with one final wash in BC150. To elute the immunoprecipitated complexes, beads were resuspended in 2x SDS loading buffer, boiled at 95⁰C for 5 min, and the supernatant collected. An appropriate amount of nuclear extract was taken as the input sample and inputs were incubated in 1x SDS loading buffer at 95⁰C for 5 min. When probing for interacting proteins of interest smaller than 50 kDa, HRP-conjugated VeriBlot secondary antibodies (Abcam) were used to avoid cross-reactivity with denatured IgG. Membranes were then imaged by chemiluminescence.

To examine the SET1A DPR/AAA mutation, HEK 293T cells were transfected with plasmids expressing WT/mutated SET1A NTD constructs using Lipofectamine 2000 according to manufacturer’s instructions. Cells were passaged the following day and allowed to grow for a further 24 hr before harvesting by trypsinisation. 600 µg nuclear extract was used as input for each IP. Extracts were diluted in nuclear extraction buffer C without salt to give a final NaCl concentration of 150 mM. Benzonase nuclease (125U) was added and extracts were incubated for 30 min at 4⁰C with gentle mixing. Samples were centrifuged at 21,000xg for 5 minutes and supernatant was used as input for IPs. 25 µl anti-FLAG M2 affinity resin (Sigma A2220) was used for each IP. Beads were washed three times in BC150 then incubated with extracts for 4 hr at 4⁰C with gentle agitation. The beads were then washed 3 times in BC150 with 0.02% NP40 and bound proteins were eluted by boiling for 5 minutes in 35 µl 2x SDS-PAGE loading dye.

### Calibrated Total RNA-sequencing (cRNA-seq)

ESCs (∼10^6^) were counted and mixed with ¼ of the number of SG4 *Drosophila* cells in PBS. RNA was extracted from cells using TRIzol reagent, according to the manufacturer’s protocol (Thermo Fisher Scientific). gDNA contamination was depleted using TURBO DNA-free Kit (Thermo Fisher Scientific) and quality of RNA was assessed using 2100 Bioanalyzer RNA 6000 Pico kit (Agilent). 900 ng RNA was depleted of rRNA using the NEBNext rRNA Depletion kit (NEB). RNA-seq libraries were prepared from an equal amount of ribo-depleted RNA using the NEBNext Ultra II Directional RNA Library Prep kit, including 2.5-4 min fragmentation at 94⁰C (NEB).

### Calibrated Transient Transcriptome-sequencing (cTT-seq)

cTT-seq was performed largely as described previously^84^. In brief, 9 million ESCs and 3 million *Drosophila* SG4 cells were labelled with 500 µM 4-thiouridine (4sU, Glentham Life Sciences) for 15 min and harvested into TRIzol reagent. 4sU-labelled mouse and *Drosophila* cells were mixed and RNA was extracted using the Direct-zol DNA/RNA Miniprep kit (Zymo Research) as per the manufacturer’s protocol. gDNA was depleted using the TURBO DNA-free Kit (Thermo Fisher Scientific). An equal quantity of RNA (60-80 µg) was taken into 100 µl nuclease free water and fragmented on ice with 20 µl 1M NaOH for 20 min. Fragmentation was stopped with 80 μl 1 M Tris, pH 6.8 and the RNA was cleaned up with Micro Bio-Spin P-30 gel columns (Biorad). RNA was biotin-labelled with 50 μl 0.1 mg/ml MTSEA biotin-XX linker (Biotium) with 3 μl biotin buffer (833 mM Tris HCl, pH 7.4, 83.3 mM EDTA) for 30 min at RT. Biotin-labelled RNA was purified with a 1:1 ratio of Phenol/Chloroform/Isoamyl alcohol (Thermo Fisher Scientific). Streptavidin pull-down was performed with the μMACS Streptavidin Kit (Miltenyi Biotec), washing the columns three times with 55°C pull-down wash buffer (100 mM Tris HCl, pH 7.4, 10 mM EDTA, 1 M NaCl and 0.1% Tween 20) and 3x RT pull down wash buffer. Biotin-labelled RNA was eluted with 100 μl elution buffer (100 mM DTT in nuclease-free water) and cleaned up with the RNeasy MinElute Cleanup kit (QIAGEN), adjusting the amount of ethanol to capture RNA < 200 nucleotides in length. RNA was quantified using the Qubit RNA HS assay kit and RNA libraries were prepared from 20-50 ng RNA with the Ultra II Directional RNA library prep kit, as per the manufacturer’s guidelines for rRNA depleted and FFPE RNA (NEB).

### Native cChIP-sequencing

Native cChIP-seq was performed as described previously^94^. In brief, 5 x 10^7^ ESCs were mixed with 2 x 10^7^ *Drosophila* SG4 cells and nuclei were released by resuspending in RSB (10 mM Tris HCl pH 8, 10 mM NaCl, 3 mM MgCl2) with 0.1% NP40. Nuclei were pelleted at 1500xg for 5 min and then washed and resuspended in 1 ml MNase digestion buffer (RSB with 0.25 M Sucrose, 3 mM CaCl2, 1x PIC). Each sample was incubated with 200 units of MNase (Fermentas) at 37⁰C for 5 minutes, with gentle inversion. Digestion was stopped by addition of 4 mM EDTA. Following centrifugation at 1500xg for 5 min, the supernatant (S1 fraction) was retained and the remaining pellet was resuspended in 300 μl nucleosome release buffer (10 mM Tris HCl pH 7.5, 10 mM NaCl, 0.2 mM EDTA, 1x PIC), rotated at 4⁰C for 1 hr and then passed five times through a 27G needle using a 1 ml syringe. Following centrifugation at 1500xg for 5 min, the supernatant (S2) was combined with S1 fraction, aliquoted, snap frozen and stored at -80°C. Digestion to predominantly mononucleosomal fragments was confirmed by agarose gel electrophoresis of purified DNA.

For each IP, 100 μl S1/S2 nucleosomes were diluted to 1 ml total volume in native ChIP incubation buffer (70 mM NaCl, 10 mM Tris HCl, pH 7.5, 2 mM MgCl2, 2 mM EDTA, 0.1 % Triton-X100, 1x PIC) and immunoprecipitated with 3 µl H3K4me3 antibody (Klose Lab) overnight at 4⁰C. IPs were all set up in duplicate for each sample. 100 μl diluted chromatin was also set aside as an input sample. Protein A agarose beads (Repligen) were blocked with 1 mg/ml BSA and 1 mg/ml yeast tRNA in native ChIP incubation buffer, overnight at 4⁰C. 40 μl slurry of pre-blocked agarose beads were used to capture antibody-bound nucleosomes at 4⁰C for 1 hr. Beads were then washed 4x with Native ChIP wash buffer (20 mM Tris HCl, pH 7.5, 2 mM EDTA, 125 mM NaCl, 0.1 % Triton-X100, 1x PIC) and 1x TE buffer, pH 8. DNA was eluted by vortexing for 30 min in elution buffer (1% SDS and 0.1 M NaHCO3) and DNA was purified using a ChIP DNA Clean & Concentrator kit (Zymo Research). For each ChIP, DNA from the matched input control (10% of the IP) was also purified. Purified DNA was analysed using ChIP-qPCR and cChIP-seq libraries for both ChIP and input samples were prepared using NEBNext Ultra II DNA Library Prep Kit for Illumina following the manufacturer’s guidelines (NEB).

### Cross-linked cChIP-sequencing

For double cross-linked T7-SET1A ChIP and ZC3H4-T7 ChIP, 5 x 10^7^ ESCs were fixed with 2 mM DSG (disuccinimidyl glutarate, Thermo Fisher Scientific) for 50 min at 25⁰C and then 1% formaldehyde (methanol-free, Thermo Fisher Scientific) for 10 min. Alternatively, for single cross-linked RNA Pol II ChIP, 5 x 10^7^ ESCs were fixed with 1% formaldehyde for 10 min at 25⁰C. Fixation was quenched using glycine added to 125 mM. Cells were then pelleted at 1000xg for 5 min and washed with PBS. Cross-linked ESCs were mixed with 1 x 10^5^ cross-linked HEK 293T/T7-SCC1 cells (1% formaldehyde, 15 min for SET1A ChIP; a gift from Martin Houlard, Nasmyth lab) or 2 x 10^6^ cross-linked HEK 293T cells (1% formaldehyde, 10 min for RNA Pol II ChIP). Chromatin was prepared by incubation in 1 ml FA-lysis buffer (50 mM HEPES pH 7.9, 150 mM NaCl, 1 mM EDTA, 0.5 mM EGTA, 0.5% NP-40, 0.1% Na-deoxycholate, 0.1% SDS, 1x PIC, 1 mM AEBSF. For RNA Pol II ChIP, EDTA concentration was increased to 2 mM and 10 mM NaF was added fresh) on ice for 10 min. Chromatin was sonicated using a BioRuptor Pico sonicator (Diagenode) at 4⁰C. Sonication was performed using 23-30 cycles of 30s on/30s off at full power, shearing genomic DNA to an average size of 0.5 kb. The sonicated material was pelleted at 20,000xg for 20 min, and the supernatant taken as sonicated chromatin.

300 μg chromatin was used per IP. Chromatin was diluted to 1 ml total volume per IP in FA-lysis buffer. An additional volume of diluted chromatin was taken to use as an input sample. Protein A agarose beads (Repligen) were blocked with 1 mg/ml BSA and 1 mg/ml yeast tRNA in 1x TE buffer at 4⁰C for 1 hr. Chromatin was pre-cleared with agarose beads (40 μl slurry beads per ChIP) at 4⁰C for 1 - 2 hr. The input sample was taken from the pre-cleared chromatin, and the remainder was immunoprecipitated overnight at 4⁰C with the appropriate amount of antibody: T7 (Cell Signalling, D9E1X, 10µl) or RNA Pol II N-terminal domain (Cell Signalling, D8L4Y, 15µl). Antibody-bound chromatin was precipitated for 3 hr at 4⁰C using 40 μl slurry of blocked Protein A agarose beads. Washes were carried out for 5 min each at 4⁰C, using FA-lysis buffer, FA-lysis buffer with 500 mM NaCl, 1x DOC buffer (10 mM Tris HCl, pH 8, 250 mM LiCl, 1 mM EDTA (2 mM EDTA for RNA Pol II ChIP), 0.5% NP-40, 0.5% Na-deoxycholate), and 2 washes with TE buffer (1x PIC and 1 mM AEBSF were added fresh to all wash buffers. 10 mM NaF was also added for RNA Pol II ChIP). DNA was eluted by vortexing for 30 min in elution buffer (1% SDS and 0.1 M NaHCO3). Cross-links were reversed for ChIPs and inputs at 65⁰C overnight with 200 mM NaCl and 2 μl RNase A (Sigma). Samples were then incubated with 20 μg Proteinase K for 1 hr at 45⁰C. DNA for ChIPs and inputs was purified using a ChIP DNA Clean & Concentrator kit (Zymo Research). Purified DNA was analysed using ChIP-qPCR. cChIP-seq libraries for both ChIP and input samples were prepared using NEBNext Ultra II DNA Library Prep Kit for Illumina following manufacturer’s guidelines (NEB).

### Massively parallel sequencing

All sequencing experiments were carried out in at least biological triplicate. Sequencing samples were indexed using NEBNext Multiplex Oligos (NEB). The average size and concentration of all sequencing libraries was analysed using a Bioanalyser High Sensitivity DNA Kit (Agilent) followed by qPCR using SensiMix SYBR Green mastermix (Bioline) and KAPA Illumina DNA standards (Roche). Libraries were sequenced using a NextSeq500 (Illumina). ChIP and TT-seq libraries were sequenced with 40 bp paired-end reads and RNA-sequencing libraries with 80 bp paired-end reads.

### Read alignment and normalisation

For cChIP-seq, paired-end reads were aligned to the concatenated mouse mm10 and spike-in genomes (mm10+dm6 for Native cChIP and mm10+hg19 for cross-linked cChIP) using Bowtie2 with the ‘-no-mixed’ and ‘-no-discordant’ options^95^. Reads that mapped more than once were discarded and PCR duplicates were removed using Sambamba^96^. For cRNA-seq and cTT-seq, reads that aligned to the mm10 and dm6 rDNA genomic sequences (GenBank: BK000964.3 and M21017.1) were first identified using Bowtie2 with ‘-very-fast’, ‘-no-mixed’ and ‘-no-discordant’ options) and discarded^95^. Unmapped reads were then aligned to the concatenated mm10 and dm6 genomes using STAR^97^. To improve mapping of intronic sequences, reads that failed to map using STAR were aligned using Bowtie2 with ‘-sensitive-local’, ‘-no-mixed’ and ‘-no-discordant’ options. Uniquely aligned reads from the last two steps were combined for further analysis and PCR duplicates were removed using Sambamba.

Sequencing datasets were calibrated to the spike-in *Drosophila* or human genomes, as described previously^94, 98-100^. For cChIP-seq, the number of mm10 reads were randomly downsampled to reflect the total number of dm6 or hg19 reads in that sample. Furthermore, to adjust for any variation in cell mixing, each sample was adjusted using the percentage of dm6 reads relative to mm10 reads in the relevant input sample. For cRNA-seq and cTT-seq, the number of mm10 reads were randomly downsampled to reflect the total number of dm6 reads in that sample. After normalisation, read coverages for individual biological replicates were compared across regions of interest using the multiBamSummary and plotCorrelation functions from deepTools (version 3.1.1)^101^. Biological replicates correlated well with each other (Pearson correlation coefficient>0.9) and were merged for subsequent analysis. Genome coverage tracks were generated using the pileup function from MACS2 for cChIP-seq and genomeCoverageBed from BEDTools (version 2.17.0) for cRNA-seq and visualised using the UCSC genome browser^102-106^. Differential genome coverage tracks (fold change of two conditions) were obtained using the bigwigCompare function from deepTools^101^.

### Peak calling and annotation

H3K4me3 peak sets were generated from each ChIP replicate using MACS2 ^103, 106^(‘BAMPE’ and ‘broad’ options specified), with a matched input sample from each biological replicate used for background normalisation. A H3K4me3 peak set comprising an overlap of the peaks called in three biological replicates in untreated samples in dTAG-SET1A/B cells were used for further analysis. TSS-associated H3K4me3 peaks were identified using BEDTools intersect^104^, only carrying forward peaks which overlapped directly with a TSS from a custom-built, non-redundant mm10 set^100^ (n=20633; TSS-associated H3K4me3 peaks n=14065). Enhancer regions were previously defined from H3K27ac cChIPseq and ATAC-seq^107^.

### Read count quantification and analysis

Differential gene expression analysis was performed as described previously^77^. In brief, mm10 read counts were obtained from individual biological replicates prior to spike-in normalisation using a SAMtools-based custom Perl script within the non-redundant mm10 gene set (n=20633). dm6 read counts were obtained from a set of unique dm6 refGene genes and were used to calculate normalisation size factors using the DESeq2 package^108^. These size factors were then applied to DESeq2 analysis of the mm10 read counts. A change was considered significant based on a threshold

---

**Extended Data Fig.1.**
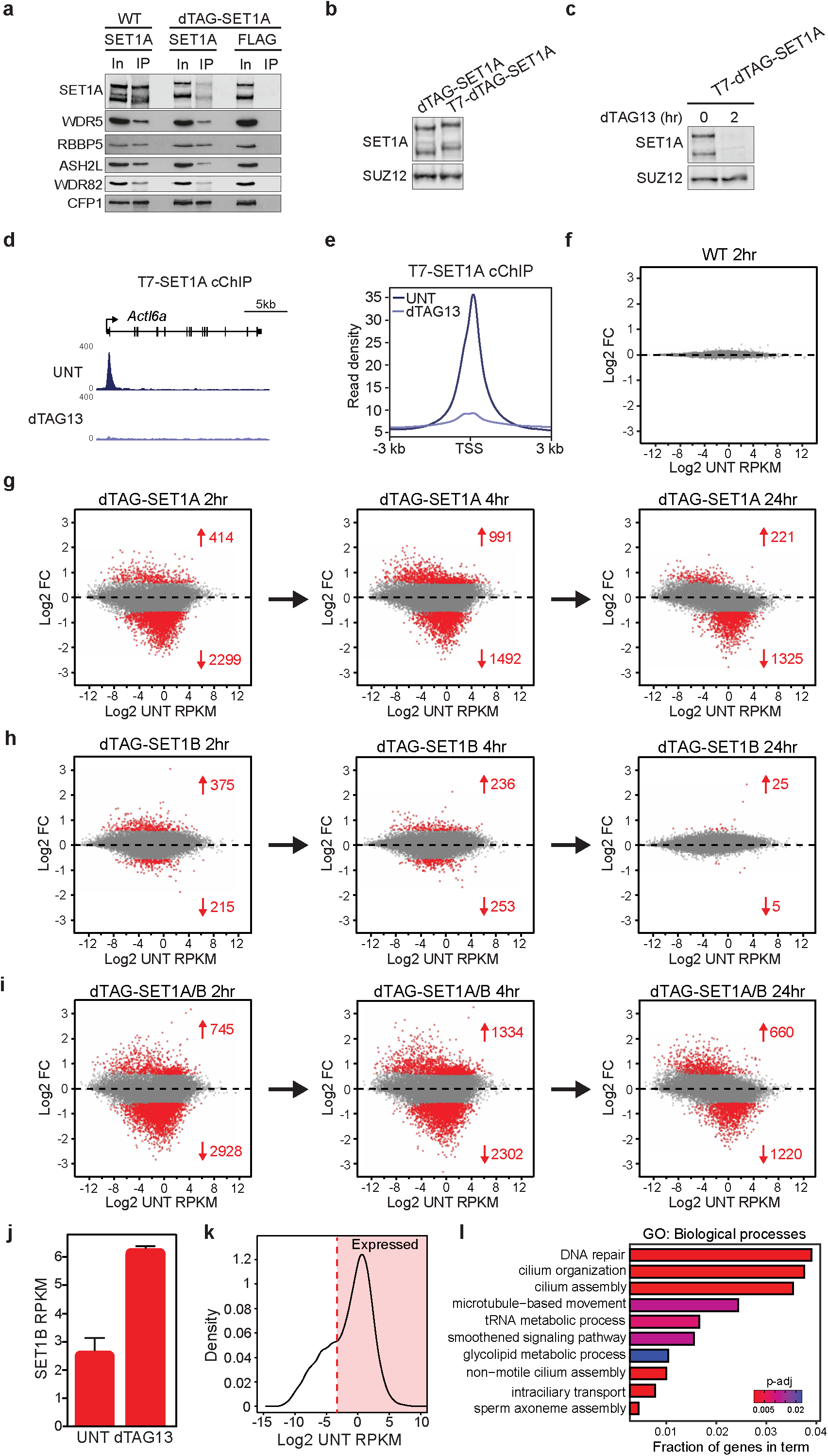
**(a)** Immunoprecipitation of SET1A from either wild type (WT) cells or dTAG-SET1A cells using a SET1A- specific antibody. This illustrates that dTAG-SET1A forms SET1 complexes normally with the other SET1 complex components WDR5, RBBP5, ASH2L, WDR82, and CFP1. The FLAG antibody immunoprecipitation is a control to indicate the specificity of the SET1A immunoprecipitations. **(b)** A western blot validating the addition of a triple T7 tag to SET1A in the dTAG-SET1A line to enable chromatin immunoprecipitation. SUZ12 functions as a loading control. **(c)** A western blot indicating that T7-dTAG-SET1A is rapidly depleted after 2 hours of dTAG13 treatment. SUZ12 functions as a loading control. **(d)** A genomic snapshot of cChIP-sequencing for T7-dTAG-SET1A before (UNT) and after 2 hours of dTAG13 treatment. SET1A signal is lost from the *Actl6a* target gene. **(e)** A metaplot of T7-dTAG-SET1A cChIP-seq signal over the transcription start site (TSS) of gene promoters before (UNT) and after dTAG13 treatment for 2 hours (n=20633). **(f)** An MA-plot showing log2 fold changes (Log2 FC) in cRNA-seq signal in wild type ESCs following dTAG13 treatment (n=20633). **(g)** An MA-plot showing log2 fold changes in cRNA-seq signal in the dTAG-SET1A line following dTAG13 treatment for 2, 4, or 24 hours (n=20633). Significant gene expression changes (p-adj<0.05 and >1.5- fold) are coloured red and the numbers of significantly changing genes are indicated. **(h)** As per (g) but for the dTAG-SET1B line. **(i)** As per (g) but for the dTAG-SET1A/B line. **(j)** A bar plot indicating the expression level (RPKM) of the *Set1b* gene before (UNT) and after 2 hours of dTAG13 treatment in the dTAG-SET1A cell line. Error bars represent SEM from three biological replicates. **(k)** A density plot of log2-transformed RPKM gene expression levels (cRNA-seq) in untreated dTAG-SET1A/B cells. The red line indicates the cut off (log2 RPKM = -3.3) that was used to separate expressed genes (n = 13028) from genes with no or very low expression. **(l)** A gene ontology (GO) analysis of genes reduced in expression after 2 hours of SET1A/B depletion. Even the most significant enriched terms account for only a very small fraction of affected genes.

**Extended Data Fig.2.**
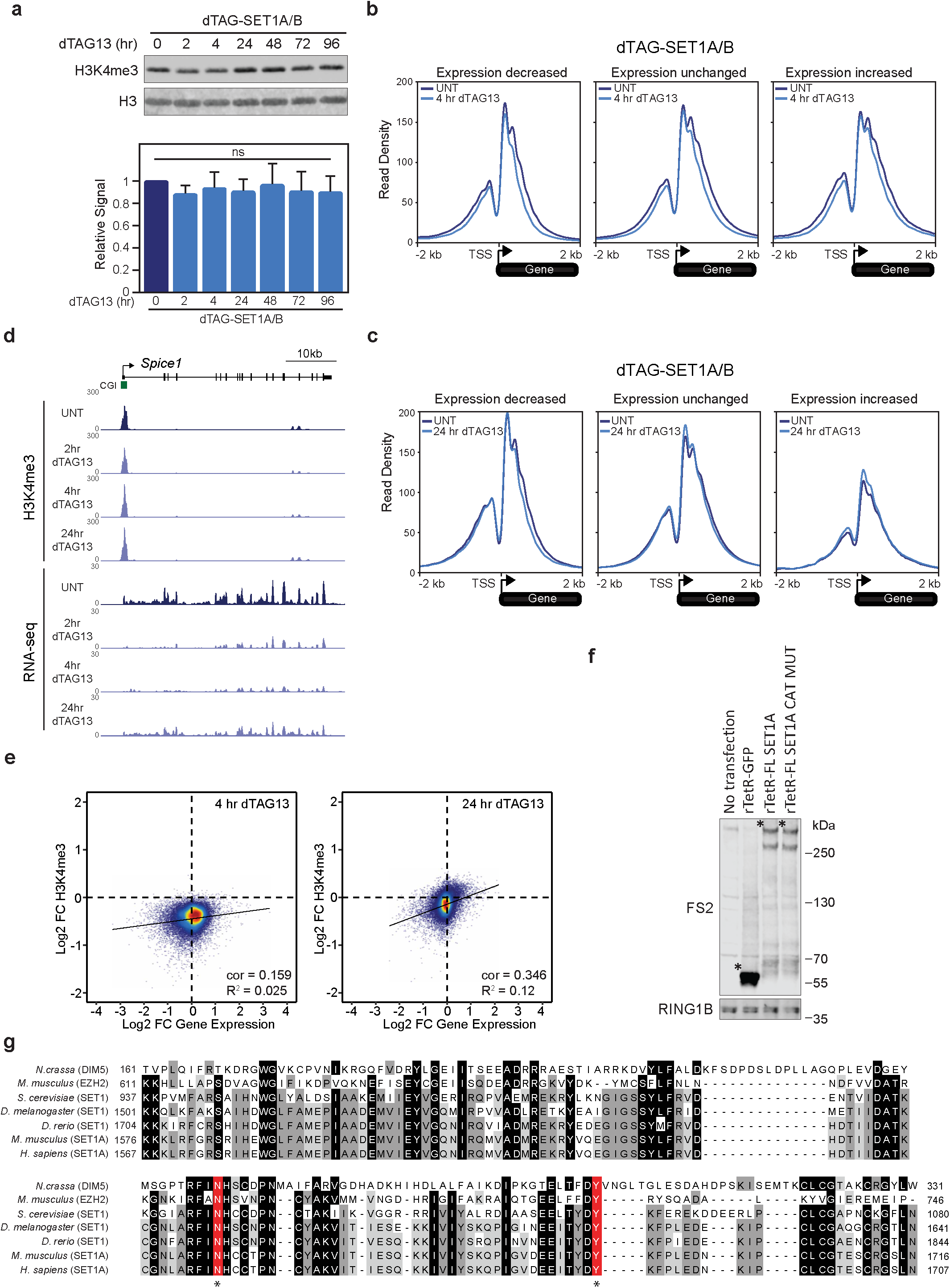
**(a)** Western blot analysis of H3K4me3 levels in the untreated dTAG-SET1A/B line and following 2, 4, 24, 48, 72, and 96 hours of dTAG13 treatment (top panels). H3 is included as a loading control. Quantitation of the alteration in H3K4me3 levels following dTAG13 treatment from 2 biological replicates. The error bars represent standard error of the mean (SEM) (bottom panel). ns indicates that the quantified changes are not significant. **(b)** Metaplot analysis of H3K4me3 cChIP-seq around the transcription start site (TSS) of genes the have reduced expression (left panel), unchanged expression (middle panel), or increased expression (right panel) in the dTAG-SET1A/B line before (dark blue lines) and after 4 hours of dTAG13 treatment (light blue lines). Only expressed genes are included (reduced expression, n=2050; unchanged, n=10081; increased expression, n=897). **(c)** As in (b) but after 24 hour of dTAG13 treatment. Only expressed genes are included (reduced expression, n=1182; unchanged, n=11567; increased expression, n=279). **(d)** A genomic snapshot comparing H3K4me3 cChIP-seq signal (top panels) and cRNA-seq (bottom panels) before and after 2,4, and 24 hours of dTAG13 treatment in the dTAG-SET1A/B line at the *Spice1* gene. **(e)** Scatter plots comparing the log2 fold change (log2FC) in H3K4me3 cChIP-seq signal and cRNA-seq signal in the dTAG-SET1A/B line before and after 4 hours (left panel) and 24 hours (right panel) of dTAG13 treatment. The correlation (cor) and R^2^ values are indicated. Only genes that have a peak of H3K4me3 in untreated cells are included (n=14065). **(f)** A western blot against the FLAG-StrepII tag (FS2) showing expression of the rTetR fusion proteins used for reporter gene expression analysis in Fig.2. In each case the * indicates the position on the blot of the FLAG-StrepII tagged rTetR fusion protein. RING1B is shown as a loading control. Molecular weight standards are shown in kilodaltons (kDa) on the right of the blot. **(g)** A multiple sequence alignment of the SET domain of various histone methyltransferases from different species. Key catalytic residues that were mutated to inactivate SET1A methyltransferase activity are shown in red. Identical residues are highlighted in black and similar residues are highlighted in grey.

**Extended Data Fig.3.**
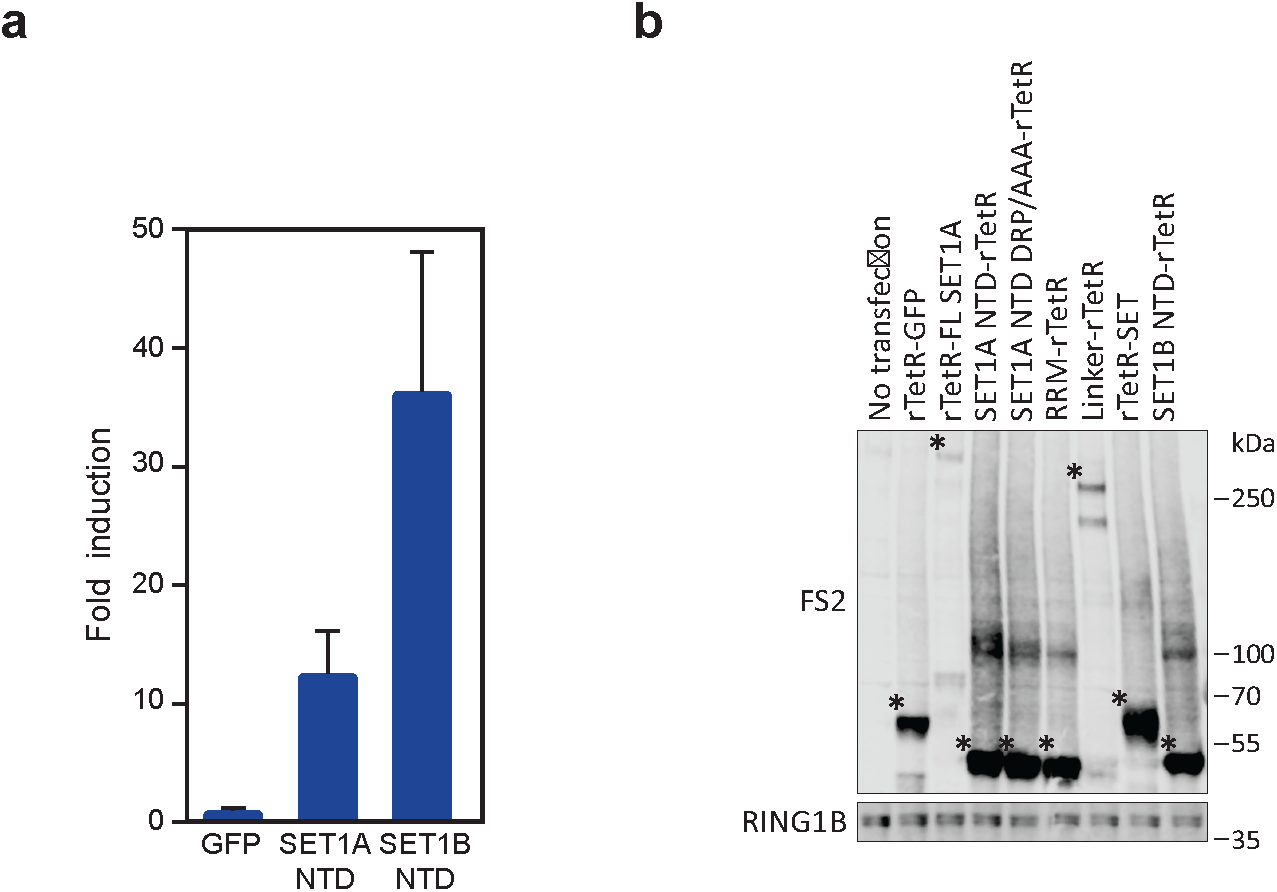
**(a)** A bar plot showing the fold induction of reporter gene expression after tethering of GFP, SET1A- NTD, and SET1B-NTD to the reporter gene. Error bars represent SEM from five biological replicates. **(b)** A western blot against the FLAG-StrepII tag (FS2) showing expression of the various rTetR fusions used for reporter gene expression analysis in Fig.3 and Extended Data Fig.3. In each case the * indicates the position on the blot of the FLAG-StrepII tagged rTetR fusion protein. RING1B is shown as a loading control. Molecular weight standards are shown in kilodaltons (kDa) on the right of the blot.

**Extended Data Fig.4.**
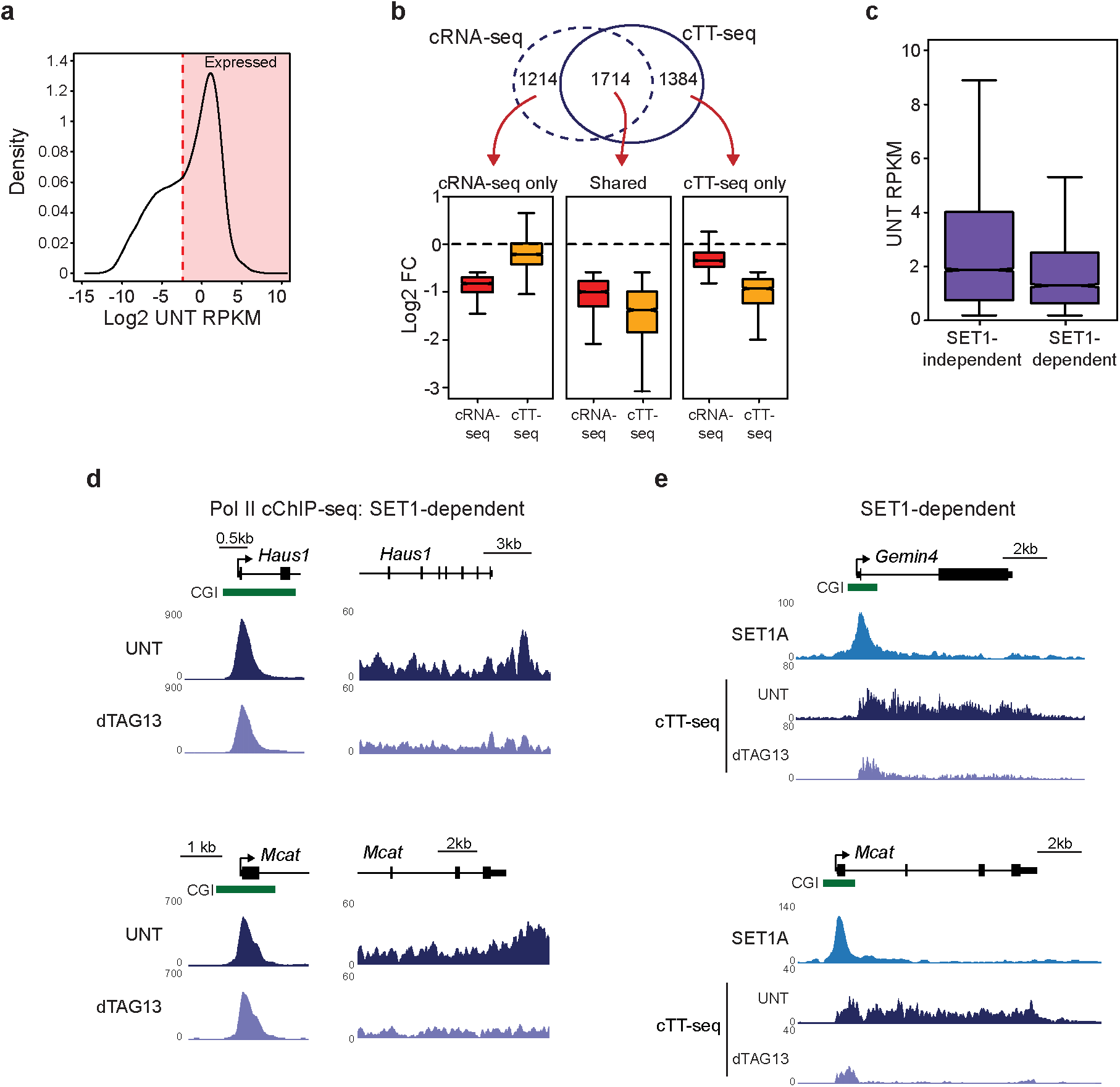
**(a)** A density plot of log 2 transformed RPKM transcription levels from cTT-seq in untreated dTAG-SET1A/B cells. The red line indicates the cut off (log2 RPKM = -2.396) that was used to separate transcribed genes (n = 11823) from genes with no or very low transcription. **(b)** A Venn diagram showing the overlap between genes significantly decreased following 2 hr SET1A/B depletion as measured by cRNA-seq or cTT-seq (top panel). Box plots of log2 fold change in gene expression, as measured using either cRNA-seq or cTT-seq, of all significantly decreased genes 2 hours after SET1A/B depletion. Genes are split into three groups, depending on whether they were significantly decreased in cRNA-seq only, cTT-seq only or in both cRNA-seq and cTT-seq (Shared). The number of genes in each subset are as per the Venn diagram. Importantly, this demonstrates that genes that only are scored as significantly reduced in expression only in either cRNA-seq or cTT-seq after SET1A/B depletion also show reductions in signal in the other assay, however, they tend to fall below the cut offs used to identify significant changes. **(c)** A box plot showing the level of transcription (RPKM) in untreated cells (UNT) for actively transcribed genes with reduced transcription when SET1A/B are depleted (SET1-dependent, n=2633) and those that are unchanged (SET1-independent, n=9151). **(d)** Genomic snapshots of RNA Pol II occupancy (Pol II cChIP-seq) at two SET1-dependent genes (*Haus1* and *Mcat*) in untreated (UNT) dTAG-SET1A/B cells (dark purple) or cells treated with dTAG13 for 2 hours (light purple). The left hand panels correspond to gene promoter occupancy and the right hand panels to gene body occupancy. **(e)** Genomic snapshots of cTT-seq signal in the dTAG-SET1A/B line at two SET1-dependent genes (*Gemin4* and *Mcat*) in untreated (UNT) cells (dark purple) or cells treated with dTAG13 for 2 hours (light purple). The location of CGIs is shown in green and SET1A cChIP-seq signal in light blue.

**Extended Data Fig.5.**
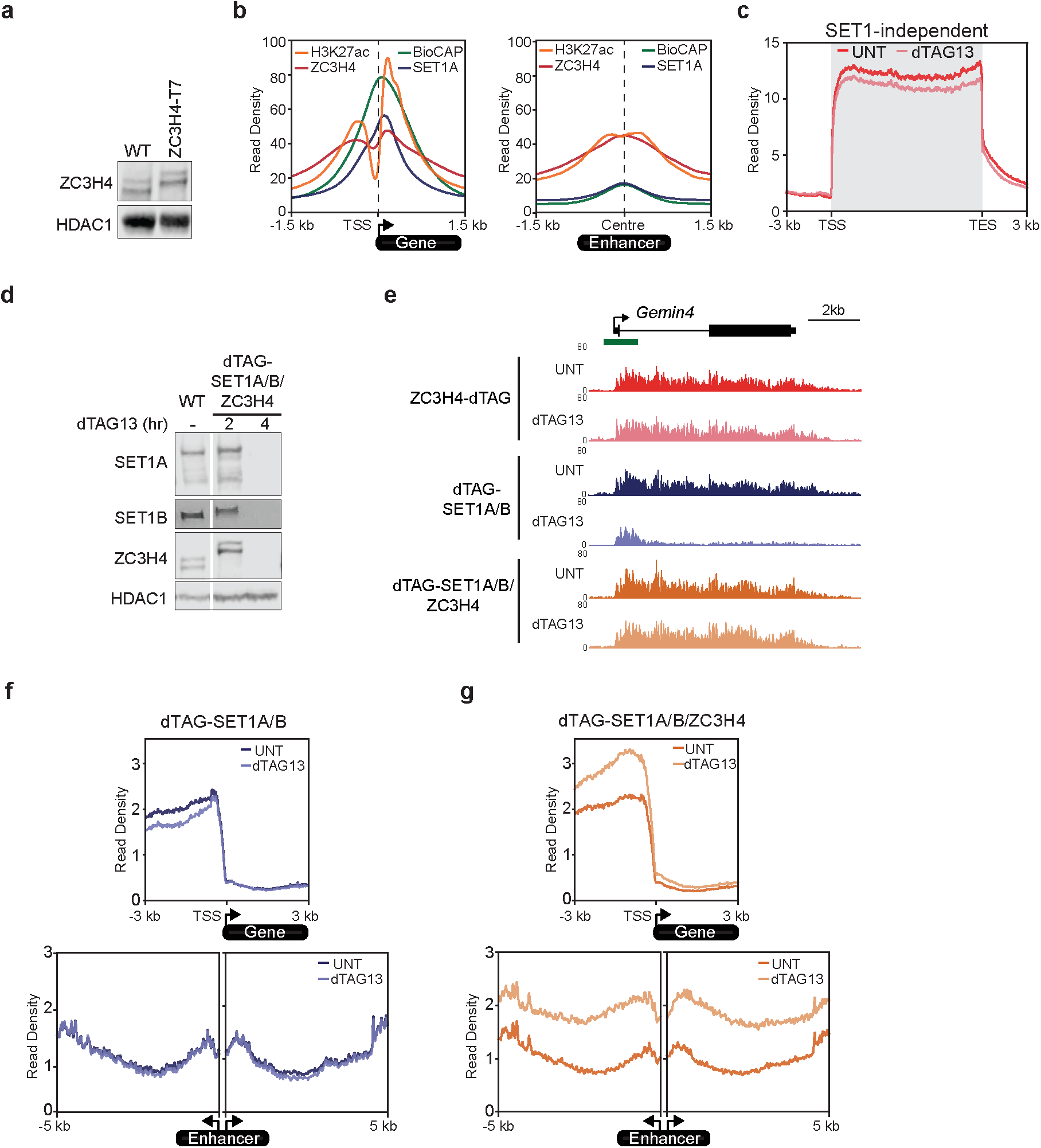
**(a)** A western blot for ZC3H4 in WT and ZC3H4-T7 cell lines. HDAC1 functions as a loading control. **(b)** Metaplot analysis of ChIP-seq signal for H3K27ac, ZC3H4, and SET1A, together with BioCAP signal, at transcribed gene promoters (left panel, n=11823) and enhancers (right panel, n=4156). BioCAP corresponds to non-methylated DNA signal in CGIs. This illustrates that ZC3H4 and H3K27 are present at promoters and enhancers. In contrast, SET1A and non-methylated DNA are enriched preferentially on the genic side of gene promoters and show little enrichment at enhancers. **(c)** Metaplot analysis of transcription (cTT-seq) in the ZC3H4-dTAG line that is either untreated (UNT) or treated with dTAG13 for 2 hours at transcribed SET1-independent genes (n=9151). **(d)** A western blot for SET1A, SET1B, and ZC3H4 in WT cells and the dTAG-SET1/ZC3H4 line. Addition of dTAG13 for 2 hours in the dTAG-SET1/ZC3H4 line leads to a simultaneous depletion of SET1A/B and ZC3H4. HDAC1 functions as a loading control. **(e)** A genomic snapshot of cTT-seq signal in the ZC3H4-dTAG, dTAG-SET1A/B and dTAG-SET1/ZC3H4 lines at a SET1-dependent gene (*Gemin4*) in untreated cells (UNT) or cells treated with dTAG13 for 2 hours. **(f)** Metaplot analysis of transcription (cTT-seq) in the dTAG-SET1A/B line either untreated or treated with dTAG13 for 2 hours. The top panel illustrates upstream antisense transcription from all gene promoters (n=20633) and the bottom panels illustrate enhancer transcription (n=4156). In both cases extragenic transcription in not significantly influenced. **(g)** Metaplot analysis of transcription (cTT-seq) in the dTAG-SET1/ZC3H4 line either untreated or treated with dTAG13 for 2 hours. The top panel illustrates upstream antisense transcription from all gene promoters (n=20633) and the bottom panels illustrate enhancer transcription (n=4156). In both cases the effects on extragenic transcription correspond to the effects caused by depletion of ZC3H4 (see Fig.5d)

## References

1. Haberle, V. & Stark, A. Eukaryotic core promoters and the functional basis of transcription initiation. Nat Rev Mol Cell Biol 19, 621–637, doi:10.1038/s41580-018-0028-8 (2018).

2. Chen, F. X., Smith, E. R. & Shilatifard, A. Born to run: control of transcription elongation by RNA polymerase II. Nat Rev Mol Cell Biol 19, 464–478, doi:10.1038/s41580-018-0010-5 (2018).

3. Core, L. & Adelman, K. Promoter-proximal pausing of RNA polymerase II: a nexus of gene regulation. Genes Dev 33, 960–982, doi:10.1101/gad.325142.119 (2019).

4. Eaton, J. D. & West, S. Termination of Transcription by RNA Polymerase II: BOOM! *Trends Genet* 36, 664–675, doi:10.1016/j.tig.2020.05.008 (2020).

5. Kamieniarz-Gdula, K. & Proudfoot, N. J. Transcriptional Control by Premature Termination: A Forgotten Mechanism. Trends Genet 35, 553–564, doi:10.1016/j.tig.2019.05.005 (2019).

6. Talbert, P. B., Meers, M. P. & Henikoff, S. Old cogs, new tricks: the evolution of gene expression in a chromatin context. Nat Rev Genet 20, 283-297, doi:10.1038/s41576-019-0105-7 (2019).

7. Illingworth, R. S. & Bird, A. P. CpG islands--’a rough guide’. FEBS Lett 583, 1713–1720, doi:10.1016/j.febslet.2009.04.012 (2009).

8. Schubeler, D. Function and information content of DNA methylation. Nature 517, 321–326, doi:10.1038/nature14192 (2015).

9. Angeloni, A. & Bogdanovic, O. Sequence determinants, function, and evolution of CpG islands. Biochem Soc Trans 49, 1109–1119, doi:10.1042/BST20200695 (2021).

10. Wu, M. et al. Molecular regulation of H3K4 trimethylation by Wdr82, a component of human Set1/COMPASS. Mol Cell Biol 28, 7337–7344, doi:10.1128/MCB.00976-08 (2008).

11. Bledau, A. S. et al. The H3K4 methyltransferase Setd1a is first required at the epiblast stage, whereas Setd1b becomes essential after gastrulation. Development 141, 1022–1035, doi:10.1242/dev.098152 (2014).

12. Fang, L. et al. H3K4 Methyltransferase Set1a Is A Key Oct4 Coactivator Essential for Generation of Oct4 Positive Inner Cell Mass. Stem Cells 34, 565–580, doi:10.1002/stem.2250 (2016).

13. van Nuland, R. et al. Quantitative dissection and stoichiometry determination of the human SET1/MLL histone methyltransferase complexes. Mol Cell Biol 33, 2067–2077, doi:10.1128/MCB.01742-12 (2013).

14. Lee, J. H. & Skalnik, D. G. CpG-binding protein (CXXC finger protein 1) is a component of the mammalian Set1 histone H3-Lys4 methyltransferase complex, the analogue of the yeast Set1/COMPASS complex. J Biol Chem 280, 41725–41731, doi:10.1074/jbc.M508312200 (2005).

15. Lee, J. H., Tate, C. M., You, J. S. & Skalnik, D. G. Identification and characterization of the human Set1B histone H3-Lys4 methyltransferase complex. J Biol Chem 282, 13419–13428, doi:10.1074/jbc.M609809200 (2007).

16. Tang, Z. et al. SET1 and p300 act synergistically, through coupled histone modifications, in transcriptional activation by p53. Cell 154, 297–310, doi:10.1016/j.cell.2013.06.027 (2013).

17. Hughes, A. L., Kelley, J. R. & Klose, R. J. Understanding the interplay between CpG island-associated gene promoters and H3K4 methylation. Biochim Biophys Acta Gene Regul Mech 1863, 194567, doi:10.1016/j.bbagrm.2020.194567 (2020).

18. Lee, J. H. & Skalnik, D. G. Wdr82 is a C-terminal domain-binding protein that recruits the Setd1A Histone H3-Lys4 methyltransferase complex to transcription start sites of transcribed human genes. Mol Cell Biol 28, 609–618, doi:10.1128/MCB.01356-07 (2008).

19. Clouaire, T. et al. Cfp1 integrates both CpG content and gene activity for accurate H3K4me3 deposition in embryonic stem cells. Genes Dev 26, 1714–1728, doi:10.1101/gad.194209.112 (2012).

20. Brown, D. A. et al. The SET1 Complex Selects Actively Transcribed Target Genes via Multivalent Interaction with CpG Island Chromatin. Cell Rep 20, 2313–2327, doi:10.1016/j.celrep.2017.08.030 (2017).

21. Mahadevan, J. & Skalnik, D. G. Efficient differentiation of murine embryonic stem cells requires the binding of CXXC finger protein 1 to DNA or methylated histone H3-Lys4. Gene 594, 1–9, doi:10.1016/j.gene.2016.08.048 (2016).

22. Eberl, H. C., Spruijt, C. G., Kelstrup, C. D., Vermeulen, M. & Mann, M. A map of general and specialized chromatin readers in mouse tissues generated by label-free interaction proteomics. Mol Cell 49, 368–378, doi:10.1016/j.molcel.2012.10.026 (2013).

23. Thomson, J. P. et al. CpG islands influence chromatin structure via the CpG-binding protein Cfp1. Nature 464, 1082–1086, doi:10.1038/nature08924 (2010).

24. Voo, K. S., Carlone, D. L., Jacobsen, B. M., Flodin, A. & Skalnik, D. G. Cloning of a mammalian transcriptional activator that binds unmethylated CpG motifs and shares a CXXC domain with DNA methyltransferase, human trithorax, and methyl-CpG binding domain protein 1. Mol Cell Biol 20, 2108–2121, doi:10.1128/mcb.20.6.2108-2121.2000 (2000).

25. Vermeulen, M. et al. Quantitative interaction proteomics and genome-wide profiling of epigenetic histone marks and their readers. Cell 142, 967–980, doi:10.1016/j.cell.2010.08.020 (2010).

26. Hung, T. et al. ING4 mediates crosstalk between histone H3 K4 trimethylation and H3 acetylation to attenuate cellular transformation. Mol Cell 33, 248–256, doi:10.1016/j.molcel.2008.12.016 (2009).

27. Saksouk, N. et al. HBO1 HAT complexes target chromatin throughout gene coding regions via multiple PHD finger interactions with histone H3 tail. Mol Cell 33, 257–265, doi:10.1016/j.molcel.2009.01.007 (2009).

28. Shi, X. et al. ING2 PHD domain links histone H3 lysine 4 methylation to active gene repression. Nature 442, 96–99, doi:10.1038/nature04835 (2006).

29. Sims, R. J., 3rd et al. Recognition of trimethylated histone H3 lysine 4 facilitates the recruitment of transcription postinitiation factors and pre-mRNA splicing. Mol Cell 28, 665–676, doi:10.1016/j.molcel.2007.11.010 (2007).

30. Taverna, S. D. et al. Yng1 PHD finger binding to H3 trimethylated at K4 promotes NuA3 HAT activity at K14 of H3 and transcription at a subset of targeted ORFs. Mol Cell 24, 785–796, doi:10.1016/j.molcel.2006.10.026 (2006).

31. Wysocka, J. et al. A PHD finger of NURF couples histone H3 lysine 4 trimethylation with chromatin remodelling. Nature 442, 86–90, doi:10.1038/nature04815 (2006).

32. Vermeulen, M. et al. Selective anchoring of TFIID to nucleosomes by trimethylation of histone H3 lysine 4. Cell 131, 58–69, doi:10.1016/j.cell.2007.08.016 (2007).

33. Sze, C. C. et al. Coordinated regulation of cellular identity-associated H3K4me3 breadth by the COMPASS family. Sci Adv 6, eaaz4764, doi:10.1126/sciadv.aaz4764 (2020).

34. Austenaa, L. M. et al. Transcription of Mammalian cis-Regulatory Elements Is Restrained by Actively Enforced Early Termination. Mol Cell 60, 460–474, doi:10.1016/j.molcel.2015.09.018 (2015).

35. Ortmann, B. M. et al. The HIF complex recruits the histone methyltransferase SET1B to activate specific hypoxia-inducible genes. Nat Genet 53, 1022–1035, doi:10.1038/s41588-021-00887-y (2021).

36. Greulich, F., Wierer, M., Mechtidou, A., Gonzalez-Garcia, O. & Uhlenhaut, N. H. The glucocorticoid receptor recruits the COMPASS complex to regulate inflammatory transcription at macrophage enhancers. Cell Rep 34, 108742, doi:10.1016/j.celrep.2021.108742 (2021).

37. Salz, T. et al. hSETD1A regulates Wnt target genes and controls tumor growth of colorectal cancer cells. Cancer Res 74, 775–786, doi:10.1158/0008-5472.CAN-13-1400 (2014).

38. Schmidt, K. et al. The H3K4 methyltransferase Setd1b is essential for hematopoietic stem and progenitor cell homeostasis in mice. Elife 7, doi:10.7554/eLife.27157 (2018).

39. Sze, C. C. et al. Histone H3K4 methylation-dependent and -independent functions of Set1A/COMPASS in embryonic stem cell self-renewal and differentiation. Genes Dev 31, 1732–1737, doi:10.1101/gad.303768.117 (2017).

40. Franks, T. M. et al. Nup98 recruits the Wdr82-Set1A/COMPASS complex to promoters to regulate H3K4 trimethylation in hematopoietic progenitor cells. Genes Dev 31, 2222–2234, doi:10.1101/gad.306753.117 (2017).

41. Clouaire, T., Webb, S. & Bird, A. Cfp1 is required for gene expression-dependent H3K4 trimethylation and H3K9 acetylation in embryonic stem cells. Genome Biol 15, 451, doi:10.1186/s13059-014-0451-x (2014).

42. Wang, L. et al. A cytoplasmic COMPASS is necessary for cell survival and triple-negative breast cancer pathogenesis by regulating metabolism. Genes Dev 31, 2056–2066, doi:10.1101/gad.306092.117 (2017).

43. Hoshii, T. et al. A Non-catalytic Function of SETD1A Regulates Cyclin K and the DNA Damage Response. Cell 172, 1007–1021 e1017, doi:10.1016/j.cell.2018.01.032 (2018).

44. Ebmeier, C. C. et al. Human TFIIH Kinase CDK7 Regulates Transcription-Associated Chromatin Modifications. Cell Rep 20, 1173–1186, doi:10.1016/j.celrep.2017.07.021 (2017).

45. Lee, J. H., You, J., Dobrota, E. & Skalnik, D. G. Identification and characterization of a novel human PP1 phosphatase complex. J Biol Chem 285, 24466–24476, doi:10.1074/jbc.M110.109801 (2010).

46. Singh, T. et al. Rare loss-of-function variants in SETD1A are associated with schizophrenia and developmental disorders. Nat Neurosci 19, 571–577, doi:10.1038/nn.4267 (2016).

47. Nagahama, K. et al. Setd1a Insufficiency in Mice Attenuates Excitatory Synaptic Function and Recapitulates Schizophrenia-Related Behavioral Abnormalities. Cell Rep 32, 108126, doi:10.1016/j.celrep.2020.108126 (2020).

48. Mukai, J. et al. Recapitulation and Reversal of Schizophrenia-Related Phenotypes in Setd1a- Deficient Mice. Neuron 104, 471–487 e412, doi:10.1016/j.neuron.2019.09.014 (2019).

49. Cenik, B. K. & Shilatifard, A. COMPASS and SWI/SNF complexes in development and disease. Nat Rev Genet 22, 38–58, doi:10.1038/s41576-020-0278-0 (2021).

50. Krebs, A. R. et al. Genome-wide Single-Molecule Footprinting Reveals High RNA Polymerase II Turnover at Paused Promoters. Mol Cell 67, 411–422 e414, doi:10.1016/j.molcel.2017.06.027 (2017).

51. Steurer, B. et al. Live-cell analysis of endogenous GFP-RPB1 uncovers rapid turnover of initiating and promoter-paused RNA Polymerase II. Proc Natl Acad Sci U S A 115, E4368–E4376, doi:10.1073/pnas.1717920115 (2018).

52. Nilson, K. A. et al. Oxidative stress rapidly stabilizes promoter-proximal paused Pol II across the human genome. Nucleic Acids Res 45, 11088–11105, doi:10.1093/nar/gkx724 (2017).

53. Erickson, B., Sheridan, R. M., Cortazar, M. & Bentley, D. L. Dynamic turnover of paused Pol II complexes at human promoters. Genes Dev 32, 1215–1225, doi:10.1101/gad.316810.118 (2018).

54. Chiu, A. C. et al. Transcriptional Pause Sites Delineate Stable Nucleosome-Associated Premature Polyadenylation Suppressed by U1 snRNP. Mol Cell 69, 648–663 e647, doi:10.1016/j.molcel.2018.01.006 (2018).

55. Almada, A. E., Wu, X., Kriz, A. J., Burge, C. B. & Sharp, P. A. Promoter directionality is controlled by U1 snRNP and polyadenylation signals. Nature 499, 360–363, doi:10.1038/nature12349 (2013).

56. Vlaming, H., Mimoso, C. A., Martin, B. J., Field, A. R. & Adelman, K. Screening thousands of transcribed coding and non-coding regions reveals sequence determinants of RNA polymerase II elongation potential. bioRxiv, 2021.2006.2001.446655, doi:10.1101/2021.06.01.446655 (2021).

57. Nojima, T. et al. Mammalian NET-Seq Reveals Genome-wide Nascent Transcription Coupled to RNA Processing. Cell 161, 526–540, doi:10.1016/j.cell.2015.03.027 (2015).

58. Schmid, M. & Jensen, T. H. Controlling nuclear RNA levels. Nat Rev Genet 19, 518-529, doi:10.1038/s41576-018-0013-2 (2018).

59. Elrod, N. D. et al. The Integrator Complex Attenuates Promoter-Proximal Transcription at Protein-Coding Genes. Mol Cell 76, 738–752 e737, doi:10.1016/j.molcel.2019.10.034 (2019).

60. Tatomer, D. C. et al. The Integrator complex cleaves nascent mRNAs to attenuate transcription. Genes Dev 33, 1525–1538, doi:10.1101/gad.330167.119 (2019).

61. Lykke-Andersen, S. et al. Integrator is a genome-wide attenuator of non-productive transcription. Mol Cell 81, 514–529 e516, doi:10.1016/j.molcel.2020.12.014 (2021).

62. Kamieniarz-Gdula, K. et al. Selective Roles of Vertebrate PCF11 in Premature and Full-Length Transcript Termination. Mol Cell 74, 158–172 e159, doi:10.1016/j.molcel.2019.01.027 (2019).

63. Li, W. et al. Systematic profiling of poly(A)+ transcripts modulated by core 3’ end processing and splicing factors reveals regulatory rules of alternative cleavage and polyadenylation. PLoS Genet 11, e1005166, doi:10.1371/journal.pgen.1005166 (2015).

64. Beckedorff, F. et al. The Human Integrator Complex Facilitates Transcriptional Elongation by Endonucleolytic Cleavage of Nascent Transcripts. Cell Rep 32, 107917, doi:10.1016/j.celrep.2020.107917 (2020).

65. Richard, P. & Manley, J. L. Transcription termination by nuclear RNA polymerases. Genes Dev 23, 1247–1269, doi:10.1101/gad.1792809 (2009).

66. Andersen, P. K., Lykke-Andersen, S. & Jensen, T. H. Promoter-proximal polyadenylation sites reduce transcription activity. Genes Dev 26, 2169–2179, doi:10.1101/gad.189126.112 (2012).

67. Ntini, E. et al. Polyadenylation site-induced decay of upstream transcripts enforces promoter directionality. Nat Struct Mol Biol 20, 923–928, doi:10.1038/nsmb.2640 (2013).

68. Austenaa, L. M. I. et al. A first exon termination checkpoint preferentially suppresses extragenic transcription. Nat Struct Mol Biol 28, 337–346, doi:10.1038/s41594-021-00572-y (2021).

69. Estell, C., Davidson, L., Steketee, P. C., Monier, A. & West, S. ZC3H4 restricts non-coding transcription in human cells. Elife 10, doi:10.7554/eLife.67305 (2021).

70. Brewer-Jensen, P. et al. Suppressor of sable [Su(s)] and Wdr82 down-regulate RNA from heat-shock-inducible repetitive elements by a mechanism that involves transcription termination. RNA 22, 139–154, doi:10.1261/rna.048819.114 (2016).

71. Berg, M. G. et al. U1 snRNP determines mRNA length and regulates isoform expression. Cell 150, 53–64, doi:10.1016/j.cell.2012.05.029 (2012).

72. Kaida, D. et al. U1 snRNP protects pre-mRNAs from premature cleavage and polyadenylation. Nature 468, 664–668, doi:10.1038/nature09479 (2010).

73. Oh, J. M. et al. U1 snRNP telescripting regulates a size-function-stratified human genome. Nat Struct Mol Biol 24, 993–999, doi:10.1038/nsmb.3473 (2017).

74. Gregersen, L. H. et al. SCAF4 and SCAF8, mRNA Anti-Terminator Proteins. Cell 177, 1797–1813 e1718, doi:10.1016/j.cell.2019.04.038 (2019).

75. Zatreanu, D. et al. Elongation Factor TFIIS Prevents Transcription Stress and R-Loop Accumulation to Maintain Genome Stability. Mol Cell 76, 57–69 e59, doi:10.1016/j.molcel.2019.07.037 (2019).

76. Nabet, B. et al. The dTAG system for immediate and target-specific protein degradation. Nat Chem Biol 14, 431–441, doi:10.1038/s41589-018-0021-8 (2018).

77. Dobrinic, P., Szczurek, A. T. & Klose, R. J. PRC1 drives Polycomb-mediated gene repression by controlling transcription initiation and burst frequency. Nat Struct Mol Biol 28, 811–824, doi:10.1038/s41594-021-00661-y (2021).

78. Raj, A., Peskin, C. S., Tranchina, D., Vargas, D. Y. & Tyagi, S. Stochastic mRNA synthesis in mammalian cells. PLoS Biol 4, e309, doi:10.1371/journal.pbio.0040309 (2006).

79. Ardehali, M. B. et al. Drosophila Set1 is the major histone H3 lysine 4 trimethyltransferase with role in transcription. EMBO J 30, 2817–2828, doi:10.1038/emboj.2011.194 (2011).

80. Hallson, G. et al. dSet1 is the main H3K4 di- and tri-methyltransferase throughout Drosophila development. Genetics 190, 91–100, doi:10.1534/genetics.111.135863 (2012).

81. Mohan, M. et al. The COMPASS family of H3K4 methylases in Drosophila. Mol Cell Biol 31, 4310–4318, doi:10.1128/MCB.06092-11 (2011).

82. Kim, J. et al. The n-SET domain of Set1 regulates H2B ubiquitylation-dependent H3K4 methylation. Mol Cell 49, 1121–1133, doi:10.1016/j.molcel.2013.01.034 (2013).

83. Hentze, M. W., Castello, A., Schwarzl, T. & Preiss, T. A brave new world of RNA-binding proteins. Nat Rev Mol Cell Biol 19, 327–341, doi:10.1038/nrm.2017.130 (2018).

84. Gregersen, L. H., Mitter, R. & Svejstrup, J. Q. Using TTchem-seq for profiling nascent transcription and measuring transcript elongation. Nat Protoc 15, 604–627, doi:10.1038/s41596-019-0262-3 (2020).

85. Schwalb, B. et al. TT-seq maps the human transient transcriptome. Science 352, 1225–1228, doi:10.1126/science.aad9841 (2016).

86. Zumer, K. et al. Two distinct mechanisms of RNA polymerase II elongation stimulation in vivo. Mol Cell 81, 3096–3109 e3098, doi:10.1016/j.molcel.2021.05.028 (2021).

87. Schulze, W. M., Stein, F., Rettel, M., Nanao, M. & Cusack, S. Structural analysis of human ARS2 as a platform for co-transcriptional RNA sorting. Nat Commun 9, 1701, doi:10.1038/s41467-018-04142-7 (2018).

88. Long, H. K. et al. Epigenetic conservation at gene regulatory elements revealed by non-methylated DNA profiling in seven vertebrates. Elife 2, e00348, doi:10.7554/eLife.00348 (2013).

89. Blackledge, N. P. et al. Variant PRC1 complex-dependent H2A ubiquitylation drives PRC2 recruitment and polycomb domain formation. Cell 157, 1445–1459, doi:10.1016/j.cell.2014.05.004 (2014).

90. Klug, M. & Rehli, M. Functional analysis of promoter CpG methylation using a CpG-free luciferase reporter vector. Epigenetics 1, 127–130, doi:10.4161/epi.1.3.3327 (2006).

91. Ran, F. A. et al. Genome engineering using the CRISPR-Cas9 system. Nat Protoc 8, 2281–2308, doi:10.1038/nprot.2013.143 (2013).

92. Faedo, A. et al. Differentiation of human telencephalic progenitor cells into MSNs by inducible expression of Gsx2 and Ebf1. Proc Natl Acad Sci U S A 114, E1234–E1242, doi:10.1073/pnas.1611473114 (2017).

93. Zhou, X., Vink, M., Klaver, B., Berkhout, B. & Das, A. T. Optimization of the Tet-On system for regulated gene expression through viral evolution. Gene Ther 13, 1382–1390, doi:10.1038/sj.gt.3302780 (2006).

94. Fursova, N. A. et al. Synergy between Variant PRC1 Complexes Defines Polycomb-Mediated Gene Repression. Mol Cell 74, 1020–1036 e1028, doi:10.1016/j.molcel.2019.03.024 (2019).

95. Langmead, B. & Salzberg, S. L. Fast gapped-read alignment with Bowtie 2. Nat Methods 9, 357–359, doi:10.1038/nmeth.1923 (2012).

96. Tarasov, A., Vilella, A. J., Cuppen, E., Nijman, I. J. & Prins, P. Sambamba: fast processing of NGS alignment formats. Bioinformatics 31, 2032–2034, doi:10.1093/bioinformatics/btv098 (2015).

97. Dobin, A. et al. STAR: ultrafast universal RNA-seq aligner. Bioinformatics 29, 15–21, doi:10.1093/bioinformatics/bts635 (2013).

98. Turberfield, A. H. et al. KDM2 proteins constrain transcription from CpG island gene promoters independently of their histone demethylase activity. Nucleic Acids Res 47, 9005–9023, doi:10.1093/nar/gkz607 (2019).

99. Blackledge, N. P. et al. PRC1 Catalytic Activity Is Central to Polycomb System Function. Mol Cell 77, 857–874 e859, doi:10.1016/j.molcel.2019.12.001 (2020).

100. Rose, N. R. et al. RYBP stimulates PRC1 to shape chromatin-based communication between Polycomb repressive complexes. Elife 5, e.18591, doi:10.7554/eLife.18591 (2016).

101. Ramirez, F., Dundar, F., Diehl, S., Gruning, B. A. & Manke, T. deepTools: a flexible platform for exploring deep-sequencing data. Nucleic Acids Res 42, W187–191, doi:10.1093/nar/gku365 (2014).

102. Kent, W. J. et al. The human genome browser at UCSC. Genome Res 12, 996–1006, doi:10.1101/gr.229102 (2002).

103. Zhang, Y. et al. Model-based analysis of ChIP-Seq (MACS). Genome Biol 9, R137, doi:10.1186/gb-2008-9-9-r137 (2008).

104. Quinlan, A. R. BEDTools: The Swiss-Army Tool for Genome Feature Analysis. Curr Protoc Bioinformatics 47, 11 12 11-34, doi:10.1002/0471250953.bi1112s47 (2014).

105. Casper, J. et al. The UCSC Genome Browser database: 2018 update. Nucleic Acids Res 46, D762–D769, doi:10.1093/nar/gkx1020 (2018).

106. Feng, J., Liu, T., Qin, B., Zhang, Y. & Liu, X. S. Identifying ChIP-seq enrichment using MACS. Nat Protoc 7, 1728–1740, doi:10.1038/nprot.2012.101 (2012).

107. Fursova, N. A. et al. BAP1 constrains pervasive H2AK119ub1 to control the transcriptional potential of the genome. Genes Dev 35, 749–770, doi:10.1101/gad.347005.120 (2021).

108. Love, M. I., Huber, W. & Anders, S. Moderated estimation of fold change and dispersion for RNA-seq data with DESeq2. Genome Biol 15, 550, doi:10.1186/s13059-014-0550-8 (2014).

